# Enzyme engineering and *in vivo* testing of a formate-reduction pathway

**DOI:** 10.1101/2021.02.15.431286

**Authors:** Jue Wang, Karl Anderson, Ellen Yang, Lian He, Mary E. Lidstrom

## Abstract

Formate is an attractive feedstock for sustainable microbial production of fuels and chemicals, but its potential is limited by the lack of efficient assimilation pathways. The reduction of formate to formaldehyde would allow efficient downstream assimilation, but no efficient enzymes are known for this transformation. To develop a 2-step formate-reduction pathway, we screened natural variants of acyl-CoA synthetase (ACS) and acylating aldehyde dehydrogenase (ACDH) for activity on one-carbon substrates and identified active and highly expressed homologs of both enzymes. We then performed directed evolution, increasing ACDH specific activity by 2.5-fold and ACS lysate activity by 5-fold. To test for *in vivo* activity of our pathway, we expressed it in a methylotroph which can natively assimilate formaldehyde. Although the enzymes were active in cell extracts, we could not detect formate assimilation into biomass, indicating that further improvement will be required for formatotrophy. Our work provides a foundation for further development of a versatile pathway for formate assimilation.

## Introduction

Population growth and climate change have created an urgent need for processes to produce more food, fuel, and chemicals while reducing CO_2_ emissions. Engineered microbes have the potential to renewably produce many useful chemicals (1). However, most commercial bioproduction uses expensive sugar feedstocks that compete with the food supply. Carbon dioxide, as a ubiquitous industrial waste and greenhouse gas, is an attractive feedstock, but CO_2_-fixing organisms are technically challenging to adapt to industrial scale. These problems can potentially be solved by bio-inorganic hybrid systems, where electricity drives catalytic production of an energy-carrying molecule used by microbes to produce value-added compounds (2, 3). Coupled to advanced photovoltaics, these systems can achieve solar-to-biomass conversion efficiencies approaching 10%, well beyond values of 3% for microalgae and 1% for plants (4).

Formate is an attractive energy carrier for a bio-inorganic system because it can be produced efficiently by electrocatalysis (5), is highly soluble in water, and provides both carbon and reducing power to microbes (2, 6). Formate can also be derived from waste biomass and fossil carbon, making it a flexible feedstock for bridging existing and future carbon economies (2).

Unfortunately, organisms that naturally consume formate are poorly suited to industrial use, and moreover, natural formate-assimilation pathways are theoretically less efficient in their consumption of ATP and reducing equivalents than rationally designed alternatives (6–8).

Recently, the first synthetic formate-assimilation pathway, the reductive glycine pathway (rGlyP), was successfully introduced into *E. coli* to support growth on formate and CO_2_ as sole carbon sources (9, 10). Although the rGlyP is energy-efficient and has great biotechnological potential, it involves a CO_2_-fixation step that requires a high ambient CO_2_ concentration in order to operate, potentially limiting its range of applications.

Several alternative formate-assimilation pathways have been proposed that could rival the efficiency of the rGlyP while not requiring CO_2_ fixation (7). These pathways all have an initial step where formate is reduced to formaldehyde, which could potentially be achieved in two enzymatic reactions via a formyl-CoA intermediate (8). For example, the ribulose monophosphate (RuMP) pathway naturally occurs in methylotrophic bacteria and assimilates formaldehyde derived from methanol oxidation (11). Assuming formate could be reduced to formaldehyde, a bacterium utilizing the RuMP pathway could easily assimilate formate as well. A second option is the rationally designed homoserine cycle, in which formaldehyde is assimilated by aldolases to generate serine or threonine, which are then assimilated by native enzymes (12). Although this pathway is not naturally occurring, its reactions can be catalyzed relatively efficiently by pre-existing *E. coli* enzymes. A few other pathways could assimilate formate in theory but will require substantial enzyme engineering to support biomass production in practice. For example, the formolase enzyme can convert formaldehyde into dihydroxyacetone (8) or glycolaldehyde (13), which can be assimilated by either natural or engineered enzymes (14). An engineered enzyme can convert formyl-CoA and formaldehyde into glycolyl-CoA and then glycolate, which can be assimilated naturally (15, 16). The common advance needed to enable all of these pathways is the reduction of formate to formaldehyde. Therefore, we sought to improve the two enzymes known to catalyze formate reduction.

## Results

### Discovery of active natural ACS variants

Previous work showed that *E. coli* acetyl-CoA synthetase (EcACS) and *L. monocytogenes* acylating acetaldehyde dehydrogenase (LmACDH) can reduce formate to formaldehyde (8). However, the wildtype enzymes tested had poor activity on the one-carbon substrates and failed to support formate reduction *in vivo* as part of the formolase pathway. To identify potential homologs of ACS with increased formyl-CoA synthetase activity, we collected 8,911 ACS sequences from UniProt and chose 41 phylogenetically diverse homologs to test experimentally (Figure S1; Materials and Methods). The chosen sequences include EcACS as well as StACSstab, a computationally stabilized variant of the *S. typhimurium* ACS well-suited to directed evolution (17). We also included ACSs from *P. aerophilum* and *K. stuttgartiensis*, which are reported to have relatively high formate activities of 27% to 65% of their acetate activities, respectively (18, 19).

We obtained the set of ACS homologs via DNA synthesis, expressed them in *E. coli*, and screened their activity in clarified *E. coli* lysates using a plate-based endpoint assay with the DTNB reagent (Materials and Methods). We performed the screens in 50mM formate, close to the K_m_ of EcACS from pilot experiments, to reveal variation in k_cat_/K_m_ across homologs. Initially we tested 11 homologs (Figure S1); using these results to highlight clades containing active variants, we then chose 30 more homologs to test. From the full set of 41 homologs, 30 had significantly higher activity than the empty vector control at a 5% false discovery rate (Figure 2A; Table S1; t-test with Benjamini-Hochberg correction), and two had higher activity than EcACS. StACSstab had lower activity than EcACS in this lysate assay, but since StACSstab is well-characterized (20), we chose to include it along with the top five ACS homologs for further analysis.

**Figure 1.**
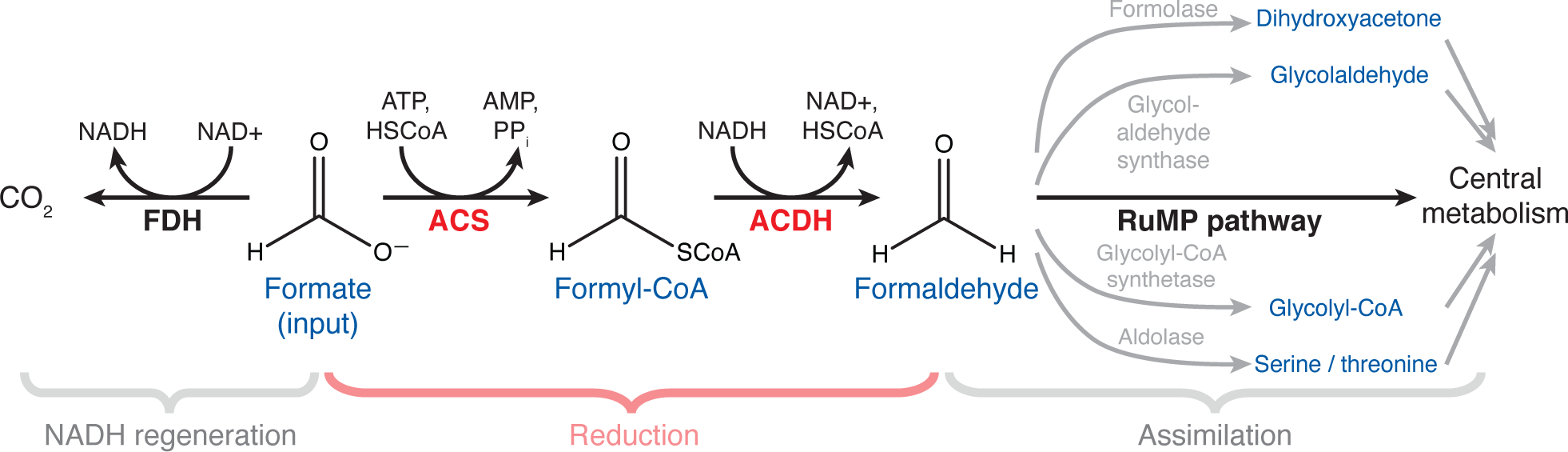
Schematic of formate reduction pathway and associated reactions. The proposed pathway reduces formate to formaldehyde via the enzymes ACS and ACDH, highlighted in red. A portion of the formate is oxidized by formate dehydrogenase (FDH) to generate the NADH needed for formyl-CoA reduction. To assimilate the formaldehyde into central metabolism and thereby support growth, the pathway is integrated into an organism which natively contains the RuMP pathway (as well as FDH). Formaldehyde could also in principle be assimilated via other pathways, such as those starting with formolase, glycolaldehyde synthase, glycolyl-CoA synthase, or a serine/threonine aldolase.

**Figure 2.**
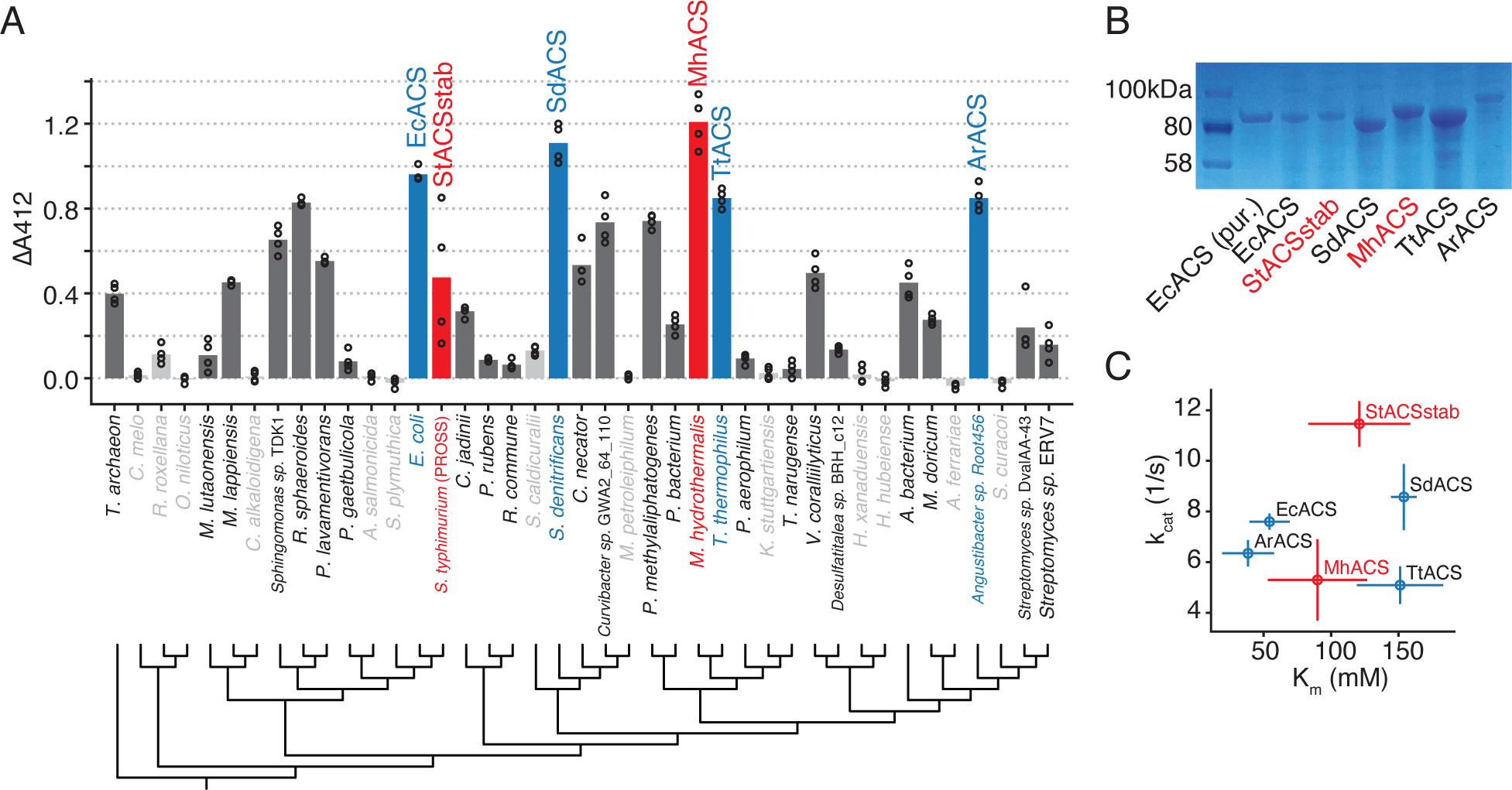
Screening natural ACS homologs identifies enzymes with formate activity. A) Lysate activity in *E. coli* for 41 ACS homologs versus their phylogeny. Activity is shown as absorbance at 412nm from the DTNB-based discontinuous assay after subtracting background (See also Figure S1; raw data in Table S1A). Circles show replicates and bars show the mean. Highlighted in color are 6 homologs chosen for purification and kinetic characterization; in red are 2 homologs chosen for directed evolution. Enzymes with statistically significant activity compared to empty vector (FDR=0.05, 2-sample t-test with Benjamini-Hochberg correction) are shown in dark gray or colored bars and black font; non-significant activity is indicated by light gray bars and font. Phylogenetic tree is a maximum-likelihood tree calculated via FastTree2 (Materials and Methods). B) SDS-PAGE on clarified lysates from 6 chosen ACS homologs. Each lane contains lysate from equal biomass. Lane 2 contains purified EcACS. C) Scatterplot of kinetic parameters k_cat_ versus K_m_ on formate of 6 chosen ACS homologs (see Figure S2 for raw kinetics data).

**Table 1.**
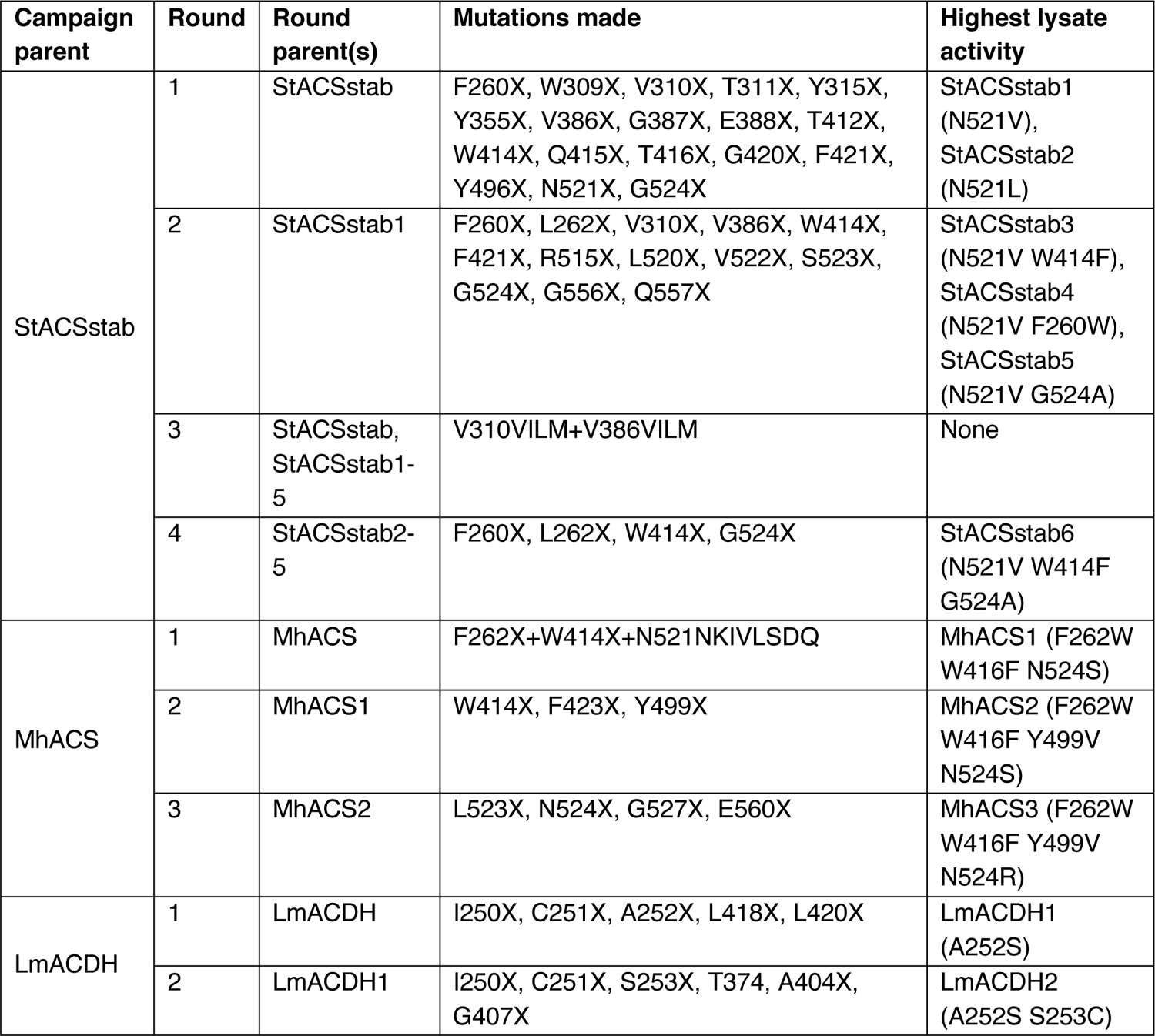
Directed evolution campaigns on StACSstab, MhACS, and LmACDH

| Campaign parent | Round | Round parent(s) | Mutations made | Highest lysate activity |
| --- | --- | --- | --- | --- |
| StACSstab | 1 | StACSstab | F260X, W309X, V310X, T311X, Y315X, Y355X, V386X, G387X, E388X, T412X, W414X, Q415X, T416X, G420X, F421X, Y496X, N521X, G524X | StACSstab1 (N521V), StACSstab2 (N521L) |
|  | 2 | StACSstab1 | F260X, L262X, V310X, V386X, W414X, F421X, R515X, L520X, V522X, S523X, G524X, G556X, Q557X | StACSstab3 (N521V W414F), StACSstab4 (N521V F260W), StACSstab5 (N521V G524A) |
|  | 3 | StACSstab, StACSstab1-5 | V310VILM+V386VILM | None |
|  | 4 | StACSstab2-5 | F260X, L262X, W414X, G524X | StACSstab6 (N521V W414F G524A) |
| MhACS | 1 | MhACS | F262X+W414X+N521NKIVLSDQ | MhACS1 (F262W W416F N524S) |
|  | 2 | MhACS1 | W414X, F423X, Y499X | MhACS2 (F262W W416F Y499V N524S) |
|  | 3 | MhACS2 | L523X, N524X, G527X, E560X | MhACS3 (F262W W416F Y499V N524R) |
| LmACDH | 1 | LmACDH | I250X, C251X, A252X, L418X, L420X | LmACDH1 (A252S) |
|  | 2 | LmACDH1 | I250X, C251X, S253X, T374, A404X, G407X | LmACDH2 (A252S S253C) |

To determine whether high lysate activity of top homologs was due to increased soluble expression, we analyzed clarified lysates by SDS-PAGE. This showed that 3 of the enzymes had much higher soluble expression than the others (SdACS, MhACS, and TtACS in Figure 2B). We then used a myokinase-coupled continuous assay to determine the kinetic parameters of purified enzymes (Figure 2C, S2; Materials and Methods). We assayed these homologs using formate as well as acetate, the likely native substrate, to determine whether any of these ACSs are already naturally biased toward one-carbon substrates. Despite its relatively low lysate activity, StACSstab had the highest k_cat_ of the 6 enzymes, on both formate and acetate (11.4 ±0.9 s^-1^ and 50.9 ± 3.1 s^-1^, respectively; Figure S4A). On the other hand, EcACS and ArACS had the lowest K_m_s (54.4 ±14.9 mM and 38.5 ±19.3 mM; Figure 2C) and highest catalytic efficiencies (k_cat_/K_m_) on formate (148 ±48 M^-1^s^-1^ and 204 ±119 M^-1^s^-1^; Figure S4A). In general, K_m_s on formate were about 3 orders of magnitude higher than on acetate. As a result, all enzymes had much lower catalytic efficiencies on formate (k_cat_/K_m_ between 50 and 200 M^-1^s^-1^) than on acetate (between 2×10^5^ and 5×10^5^ M^-1^s^-1^) (Figure 3A), although there is some variation in this specificity ratio (Figure S4B). The measured k_cat_values were also generally lower for formate than for acetate, although usually by less than one order of magnitude. In one case, SdACS actually has higher formate k_cat_ (8.6 ±1.3 s^-1^) than acetate k_cat_ (7.0 ±0.4 s^-1^).

**Figure 3.**
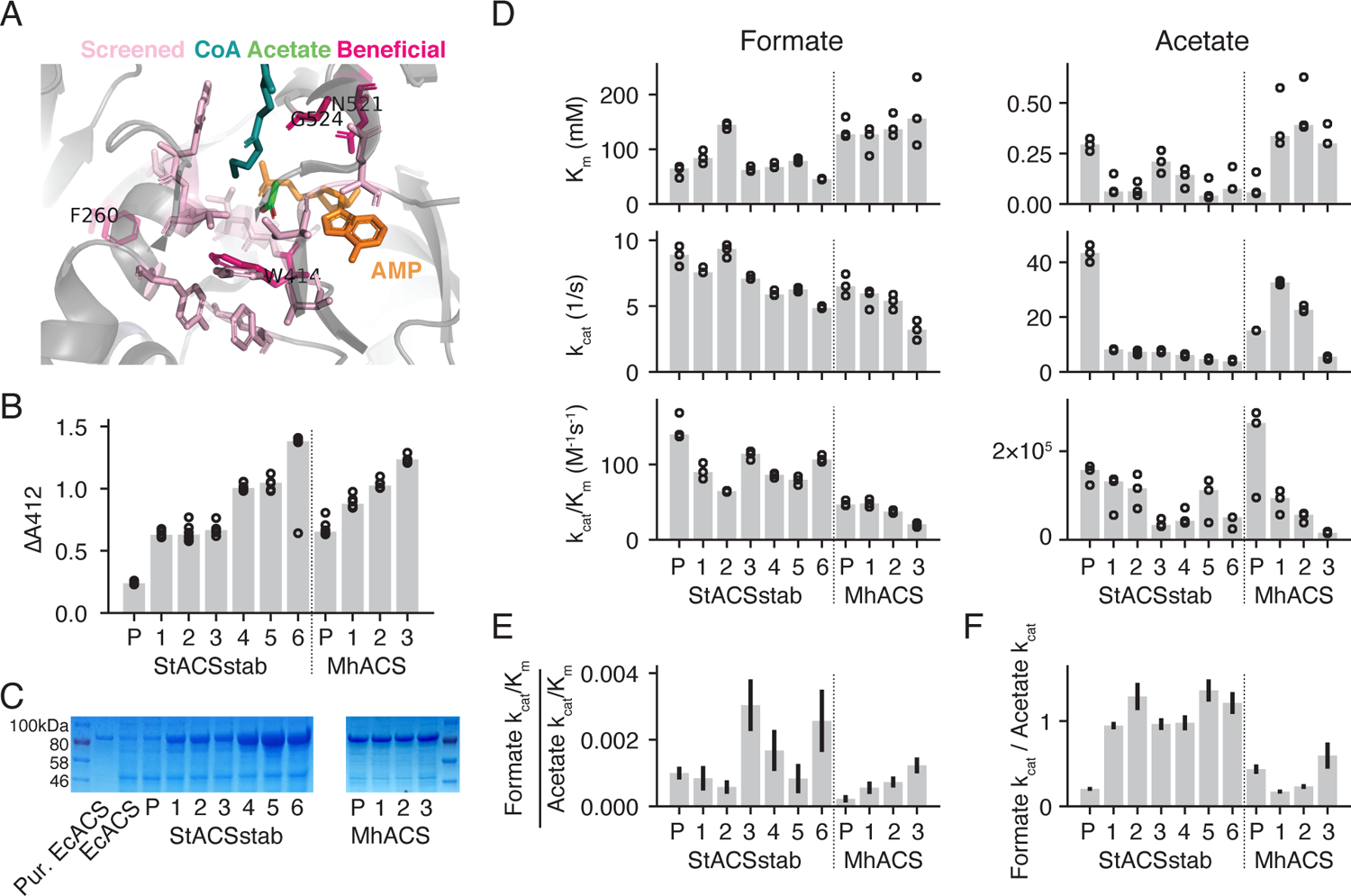
Directed evolution of ACS improves expression and specificity, but not specific activity. A) Structure of *S. typhimurium* ACS (PDB: 2p2f) with residues chosen for site-saturating mutagenesis highlighted in light pink. Residues that yielded beneficial mutations that were kept in the evolved variants are highlighted in dark pink. B) Background-subtracted lysate activity in *E. coli* using the discontinuous assay (Materials and Methods) on evolved variants of StACSstab and MhACS. All enzyme variants shown were isolated in host strain BL21*(DE3), except MhACS3, which was isolated in NovaBlue(DE3) (Figure S6). Circles show 3 replicates and bars show the median. “P” indicates parental or wildtype enzyme, and numbered variants correspond to mutants listed in Table 1. C) SDS-PAGE of clarified lysates of expression cultures of each evolved variant. Each lane contains lysates from equal biomass. D) Km, *k_cat_*, and *k_cat_*/*k_m_* for formate and acetate of evolved variants. Circles show 3 replicates and bars show the median. These parameters are also listed in Table S2. E) Ratios of formate *k_cat_*/*k_m_* to acetate *k_cat_*/*k_m_*. F) Ratios of formate *k_cat_* to acetate *k_cat_*. Error bars represent s.d. of the ratios estimated using the replicate data.

### Directed evolution of ACS

Given that even the most active of these ACS homologs is still 2-3 orders of magnitude less efficient on formate than on acetate, we performed directed evolution to increase the formate activity of ACS. No single homolog simultaneously had the highest activity, expression, and specificity, so we chose two parent enzymes: StACSstab because it had the highest k_cat_, and MhACS (from *M. hydrothermalis*) because it had the highest lysate activity. Both are also likely tolerant to mutation, since StACSstab is computationally stabilized and MhACS comes from a thermophile (21).

We took a semi-rational approach to engineer StACSstab using a published crystal structure of the wildtype StACS (PDB: 2p2f). StACSstab has 46 mutations relative to StACS (93% amino-acid identity), but these mutations were designed to avoid perturbing the structure of the active site (17). Therefore, we used the StACS structure and previous mutagenesis studies to choose a set of 18 residues lining the active-site pocket near the acyl moiety for mutagenesis (Figure 3A, Table 1) (Materials and Methods) (20,22,23). We screened lysates of single-site-saturating libraries of these 18 positions in StACS using a continuous assay in 50mM formate, followed by a secondary screen with plate-based purification (Figure S5; Materials and Methods). We identified 2 mutations, N521V (StACSstab1) and N521L (StACSstab2) that increased lysate activity by almost 3-fold (Figure 3B). Then, using StACSstab1 as a parent, we mutated positions that were beneficial in round 1 as well as new positions that were structurally proximal to N521, screening using the discontinuous assay. We isolated N521V W414F, N521V F260W, and N521V G524A (StACSstab3, StACSstab4, and StACSstab5, respectively) as variants with further improved lysate activity (Figure 3B). Previous work found that V310 and V387, which line the acyl-binding pocket in ACS, play a strong role in controlling substrate specificity (20, 22).

Therefore, in a third round of evolution, we combinatorially mutated these 2 positions to each of 3 larger hydrophobic amino acids. However, this failed to generate any improved variants (Table 1). In a final round, we combined the mutations discovered in round 2 and identified N521V W414F G524A (StACSstab6) as the most-improved candidates (Figure 3B). From StACSstab to StACSstab6, lysate activity increased by 5.8-fold (ratio of median of 4 replicates in Figure 3B).

All mutant libraries were from single-site-saturating mutagenesis (e.g. “F260X, W309X” denotes 2 libraries, of which one contains F260 mutated to all 20 substitutions, the other W309 mutated to all 20 substitutions), unless specific subsets of substitutions are indicated. Plus sign “+” indicates a combinatorial library at multiple positions. Screening data for a subset of evolution rounds is shown in the supplemental figures.

To determine whether increases in lysate activity translated to increases in specific activity, we purified the ACS variants and measured their kinetic parameters. The initial variants StACSstab1 and StACSstab2 have similar *k_cat_* on formate to the parent enzyme (7-10 s^-1^), but have 5.3-fold and 6.0-fold lower *k_cat_* on acetate (Figure 3D), respectively. This is accompanied by higher soluble expression (Figure 3C), suggesting that high levels of native (acetate) activity may be toxic and prevents high expression of parental StACSstab. Subsequent variants StACSstab2 through StACSstab6 continued to increase in soluble expression as well as formate specificity. The final variant StACSstab6 had a ratio of formate to acetate *k_cat_*/*k_m_* 2.6-fold higher than that of StACSstab. The formate to acetate *k_cat_* ratio increased even more, by 5.9-fold, between these enzymes.

Although directed evolution increased expression and formate specificity, it did not increase formate activity. In fact, catalytic activity decreased modestly over the course of evolution. The final mutant, StACSstab6, has a formate *k_cat_* of 4.9 ±0.1 s^-1^ and *k_cat_*/*k_m_* of 108 ±5 M^-1^s^-1^, 45% and 28% lower, respectively, than StACSstab’s *k_cat_* of 8.8 ±0.8 s^-1^ and *k_cat_*/*k_m_* of 149 ±18 M^-1^s^-1^ Figure 3D, Table S2). Normalizing the relative change in lysate activity to that of *k_cat_*/*k_m_*, we estimate the change in functional expression to be 8-fold, consistent with the qualitative increase in band intensity on the SDS-PAGE gel.

To engineer MhACS, we mutated a small set of positions corresponding to those in StACSstab that yielded beneficial mutations (Table 1). We first screened a combinatorial library with mutations at positions F262, W416, N524, and G527 (corresponding to StACSstab F260, W414, N521, and G524), which yielded a mutant F262Y W416F N524S (MhACS1) with improved lysate activity. Using MhACS1 as a parent, we then screened single-site-saturating libraries and discovered an improved variant with additional mutation Y499V (MhACS2). At this point, screening additional site-saturating libraries MhACS2 failed to yield improved mutants. Given the correlation between higher StACSstab expression and lower ACS activity, we hypothesized that an *E. coli* host strain with lower basal expression may increase our chances of isolating more improved mutants. Therefore, we switched from the BL21*(DE3) host strain to NovaBlue(DE3), which has the stronger LacI^q^ repressor (Figure S6). Starting with MhACS2 and screening site-saturating libraries, we obtained a mutant with S524R (MhACS3; N524R relative to MhACS) with 1.9-fold higher lysate activity than the MhACS parent (Figure 3B).

As with StACSstab, increased lysate activity of MhACS mutants did not translate to increased specific activity, but rather a decrease in formate *k_cat_* and *k_cat_*/*k_m_* over the course of evolution (Figure 3D). Formate *k_m_* also did not change appreciably, staying around 150 mM in all variants (Figure 3D, Table S2). Like StACSstab, the *k_cat_*/*k_m_* for acetate decreased in successive rounds of mutation. Thus, the ratio of formate to acetate *k_cat_*/*k_m_* increased by 5.5-fold from MhACS to MhACS3, although its absolute value is lower for MhACS3 than for StACSstab6. Interestingly, the decrease in acetate *k_cat_*/*k_m_* in MhACS3 was due to a combination of increased *k_m_* and decreased *k_cat_* contributed by different mutations. By contrast, in the StACSstab evolutionary trajectory, the major change was a drastic decrease in *k_cat_* caused by the initial N521V mutation, which was actually negated somewhat by a decrease in *k_m_*. Normalizing the increase in lysate activity from MhACS to MhACS3 by the 59% decrease in formate *k_cat_*/*k_m_*, we find that functional expression of MhACS3 is 4.6-fold higher than that of MhACS. This is surprising given the roughly similar apparent expression of all MhACS variants, perhaps suggesting a change in the active fraction of expressed protein.

### Discovery and improvement of ACDH

Previous work used *Listeria monocytogenes* (LmACDH) for formate reduction because it was the most active of 5 homologs tested (8). We sought to identify additional active variants by a two-pronged strategy of homolog screening and directed evolution. Although formyl-CoA is the desired substrate of ACDH, it is not commercially available and has a very short half-life(15). Therefore, we screened ACDHs using formate as a substrate instead, including ACS as a coupling enzyme to generate formyl-CoA in the reaction (Figure S7A). This does not allow quantitative estimation of the *k_m_* of ACDH for formyl-CoA but is sufficient to determine the relative activities of ACDH variants.

We analyzed all available ACDH homologs in UniProt and BRENDA, and chose 46 for gene synthesis and testing. Alignment and clustering of ACDHs revealed two divergent clades with roughly equal numbers of sequences (Figure 4). One clade contained members such as *E. coli* MhpF, which natively operates as a complex with an aldolase (24). EcMhpF was previously shown to have low activity compared to LmACDH(8), suggesting difficulties in expressing the monomer form. Therefore, we avoided members of the MhpF-like clade and focused instead on the clade containing *E. coli* AdhE, LmACDH, and bacterial-microcompartment-associated enzymes such as EutE (25).

**Figure 4.**
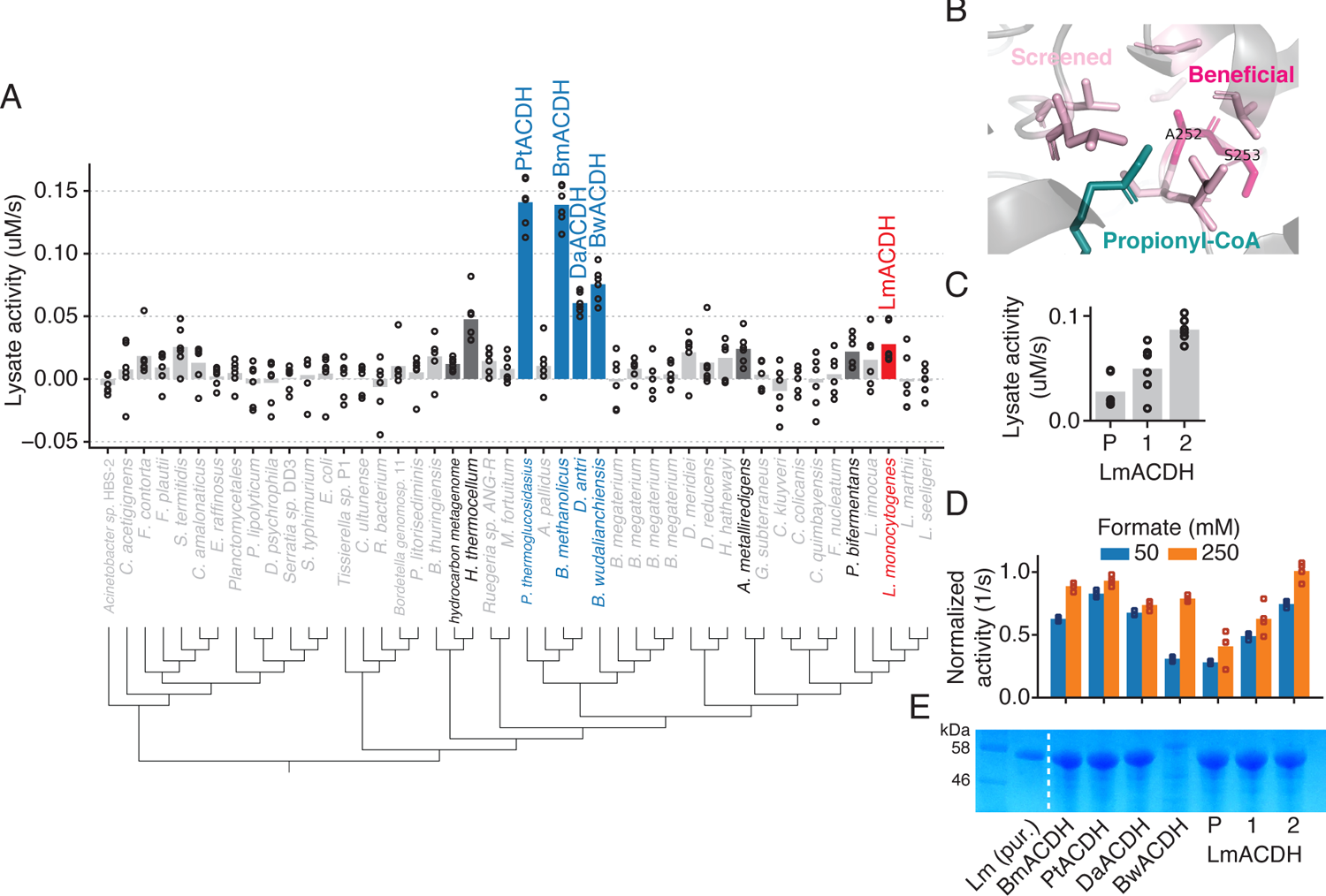
Homolog screening and directed evolution of ACDH enzymes. A) Lysate activity of ACDH homologs. Dots show technical replicates from 2 independent expression strain transformants; bars show mean of all replicates. Enzymes with statistically significant activity compared to empty vector (FDR=0.05, 2-sample t-test with Benjamini-Hochberg correction) are shown in dark gray or colored bars and black font; non-significant activity is indicated by light gray bars and font. Blue indicates enzymes chosen for followup characterization and red indicates enzyme used for directed evolution. Raw data in Table S1B. B) Structure of *L. monocytogenes* ACDH (3k9d, propionyl-CoA from 5jfn) with residues chosen for site-saturating mutagenesis shown in light pink. Residues that yielded beneficial mutations that were kept in the evolved variants are highlighted in dark pink. C) Lysate activity of evolved LmACDH variants. “P” indicates parental or wildtype enzyme, and numbered variants correspond to mutants listed in Table 1. Circles represent 3 technical replicates and bars show the mean. D) Activity of selected homologs and evolved LmACDH variants, normalized to enzyme concentration. Shares x-axis labels with panel E. E) SDS-PAGE of clarified lysates of *E. coli* strains expressing ACDH variants. Each lane contains lysate from equal biomass. Lane 2 contains purified LmACDH.

We screened clarified lysates of the ACDH homologs for the ability to oxidize NADH in the presence of ACS, ATP, CoA, and 50mM formate. We initially screened 9 ACDH homologs and then used those results to choose 37 more (Figure S1). From the full set of 46 homologs, we found 9 with activity significantly higher than empty vector at a 5% false discovery rate (Figure 4A, S8A; t-test with Benjamini-Hochberg correction). Five homologs had higher activity than LmACDH (Figure 4A, colored bars). We then purified the 4 homologs with highest lysate activity as well as LmACDH and assayed them in 50mM and 250mM formate with excess ACS. Unlike in our ACS screen, all the ACDH homologs with higher lysate activity than LmACDH also had higher activity after normalizing by enzyme concentration, with PtACDH having the highest activity in both assays (normalized activity of 0.93 ± 0.04 s^-1^ in 250mM formate; Figure 4D, S8B). All homologs were more active in 250mM formate than in 50mM. Their relative rankings were unchanged by formate concentration, except for BwACDH, which is the least active in 50mM formate but among the most active homologs at 250mM. This suggests that it has a higher *k_m_* for formyl-CoA than other homologs.

To engineer LmACDH, we used its crystal structure 3k9d along with a propionyl-CoA substrate superimposed from a related structure 5jfn to choose positions close to the acyl moiety for mutagenesis. We screened site-saturating libraries at 5 positions and identified a mutant A252S (LmACDH1) with increased activity (Table 1). We then screened some of the same positions on top of the LmACDH1 background, as well as additional residues close to A252 in the structure, and found A252S S253C (LmACDH2) to have even higher activity (1.00 ± 0.07 s^-1^ at 250mM formate, or 2.5-fold higher than LmACDH; Figure 4D). In fact, LmACDH2 has slightly higher activity than PtACDH, the best homolog we discovered.

### Expression of pathway in *M. flagellatus* KT

Having identified ACSs with improved expression and ACDHs with increased activity, we next asked whether these enzymes could support formate reduction *in vivo*. Methylotrophic bacteria are able to assimilate formaldehyde as an intermediate of methanol, and those that do this via the RuMP pathway cannot natively assimilate formate. Therefore, if we introduced ACS and ACDH activities into an RuMP methylotroph (which also had an NADH-producing FDH), this would in principle confer partial or complete formatotrophy. In practice, our enzymes are likely too inefficient to support growth, but even a low flux from formate into biomass could potentially be used to select for further enzyme improvements.

We chose the betaproteobacterium *Methylobacillus flagellatus* KT to express the pathway because it assimilates methanol via the RuMP pathway, grows robustly under standard laboratory conditions, and is amenable to genetic manipulation (26). We first tested a panel of promoters for their ability to drive high constitutive expression of a red fluorescent protein reporter from a IncP-based broad-host-range plasmid in *M. flagellatus* KT (27) (Figure S9).

Based on this, we chose to use the native promoters Phps and PmxaF to drive ACS and ACDH expression, respectively, from the plasmid. We cloned a panel of expression vectors containing different ACSs coexpressed with the same ACDH, or vice versa, and conjugated them into *M. flagellatus* KT (Figure 5A; Materials and Methods).

**Figure 5.**
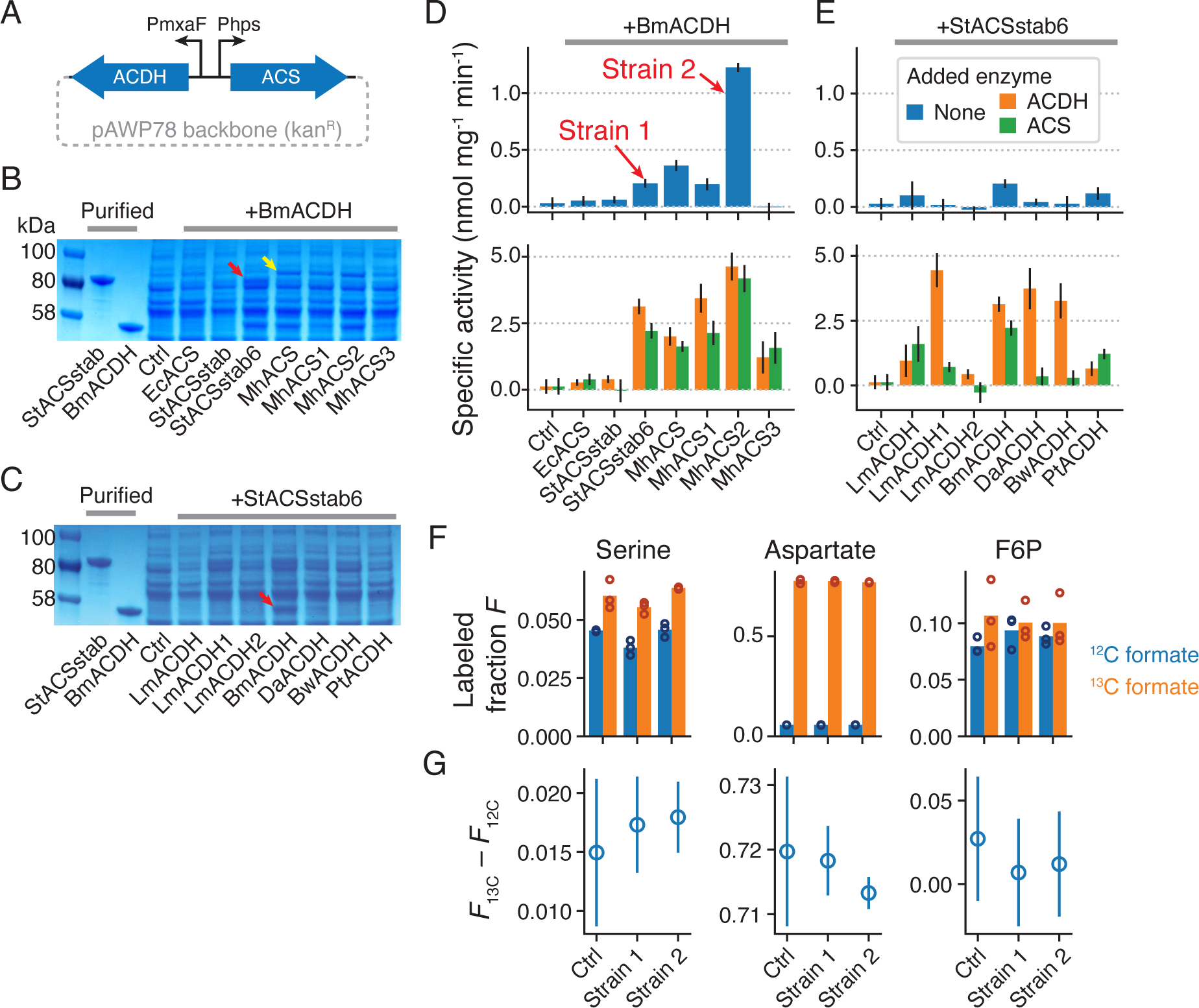
ACS and ACDH are expressed and active in *M. flagellatus* KT lysates but do not assimilate formate *in vivo*. A) Schematic of vector used to express ACS and ACDH variants in *M. flagellatus* KT. PmxaF and Phps are native promoters (see also Figure S9). B) SDS-PAGE of clarified lysates from *M. flagellatus* KT strains expressing different ACSs, as well as BmACDH, from the expression vector. Control strain (“Ctrl”) contains the same vector, but with a pJ23101-dTomato insert instead of ACS/ACDH enzymes. Red arrow: gel band for StACSstab6; yellow arrow: gel band for MhACS. C) Same as (B), but for different ACDHs co-expressed with StACSstab6. Red arrow: gel band for BmACDH. D) Specific activity of *M. flagellatus* KT lysates for formaldehyde production from 300mM formate, for various ACSs co-expressed with BmACDH (mean and s.d. from 3 technical replicates; see Materials and Methods, Figure S10). Results for cell lysates only (blue bars) or with 4µM purified ACDH (orange bars) or ACS (green bars) added. E) Same as (D), but for various ACDHs co-expressed with StACSstab6. “Strain 1” and “Strain 2” were chosen for ^13^C labeling. F) Fraction of ^13^C-labeled proteinogenic serine, aspartate, or fructose-6-phosphate (F6P) from cells grown in methanol + 200mM ^12^C or ^13^C formate. 3 biological replicates of the control strain and 2 different pathway variants (“Strain 1” and “Strain 2” from (D)) were assayed. Bars show the mean. G) The difference in ^13^C-labeled fraction between ^13^C-formate-grown cells and ^12^C-formate-grown cells (mean and s.d., n=3). If ^13^C formate were being assimilated by the pathway, then the pathway-containing strains should have a higher value of this difference than the control strain.

To test for expression and activity of our enzymes in *M. flagellatus* KT, we analyzed clarified lysates by SDS-PAGE and Nash assay (28), which measures formaldehyde production (Figure S10; Materials and methods). Formaldehyde is only produced from formate if both ACS and ACDH are active, so we assayed lysates with and without an added excess of purified ACS or ACDH to detect activity of each enzyme individually (Figure 5D,E). This also has a side benefit of boosting the sensitivity of the assay. Independent *M. flagellatus* KT transconjugants varied in phenotype (growth rates, enzyme activities), so for each vector we screened multiple transconjugants and picked the one with the highest enzyme activity for further characterization.

We found that StACSstab6 and all MhACS variants, but not EcACS or StACSstab, had a visible band in SDS-PAGE when expressed in *M. flagellatus* KT. MhACS has a higher-molecular-weight band than StACSstab6, reflecting their predicted molecular weights (74 and 72 kDa, respectively). Consistent with the SDS-PAGE, the Nash assay only showed ACS activity in StACSstab6 and MhACS variants (orange bars in Figure 5D, lower panel). Across ACDHs, only BmACDH had visible SDS-PAGE expression. It also had the highest ACDH activity by Nash assay, although LmACDH, LmACDH1, and PtACDH also had above-background activities (green bars in Figure 5E, bottom panel). The strains with the highest lysate activity without any added enzymes are those containing BmACDH and either StACSstab6, MhACS, MhACS1, or MhACS2. Notably, neither EcACS nor the LmACDH variants, which were used in the previous version of the pathway (8), were well-expressed in *M. flagellatus* KT. We chose the strains containing StACSstab6/BmACDH (“Strain 1”), which had the 3rd-highest activity (0.20 ± 0.04 nmol mg^-1^ min^-1^; Figure 5D), and MhACS2/BmACDH (“Strain 2”), which had the highest activity (1.22 ± 0.04 nmol mg^-1^ min^-1^), for further characterization.

Some strains with the same ACDH or ACS differed in their apparent activities for that enzyme in the Nash assay. This could be due to cryptic variation between transconjugants, or some interaction between the divergent Phps and PmxaF promoters. The latter might explain why, for example, across the MhACS variants in Figure 5D, ACDH activity seems to vary and correlate with ACS activity even though the ACDH enzyme is the same. We verified that the sequence of the promoters and enzyme genes on the expression vector are as expected in every strain. Therefore, any cryptic genetic variation would have to be in the genome of the host strain.

### Test for formate assimilation in *M. flagellatus* KT

Given the observed activity of ACS and ACDH in *M. flagellatus* KT lysates, we next tested for assimilation of formate into biomass. To do this, we cultured Strain 1 and Strain 2 in ^12^C- or ^13^C-formate and monitored ^13^C labeling of metabolites via liquid chromatography and mass spectrometry (LC-MS). Because *M. flagellatus* KT contains formate dehydrogenases capable of oxidizing formate to CO2, which can potentially be reassimilated via carboxylation, we also assayed a control strain containing an dTomato-expressing vector. If formate is being assimilated into biomass via our pathway, we should observe more ^13^C labeling in central carbon metabolites and proteins in a pathway-containing strain relative to the control strain, and only when labeled formate is provided.

First, we analyzed proteinogenic amino acids as an indicator of overall incorporation of formate into biomass. We inoculated control and pathway strains into MM2 medium with 0.2% ^12^C methanol and 200mM ^12^C or ^13^C-formate, harvested the saturated cultures, and acid-hydrolyzed the biomass for LC-MS. We found almost no labeling of more than one carbon atom across the amino acids examined, so we used total ^13^C-labeled fraction, or 1 – unlabeled fraction, as a simple metric for the degree of labeling (Figure 5F, S11). Serine, whose carbon atoms are derived from pyruvate and thus immediately downstream of the RuMP pathway, displayed a background ^13^C-labeled fraction of about 4% in ^12^C-formate across all strains, but a 1.5 - 2% increase in labeling in ^13^C-formate (Figure 5F). However, this increase occurred in both control and pathway strains and was similar in magnitude (Figure 5G), indicating that the extra labeling is not due to formate assimilation via our pathway. Aspartate had much higher labeling in ^13^C-formate than in ^12^C-formate, although the differential labeling was again the same in all strains (Figure 5F,G, middle panel). Since aspartate is derived from oxaloacetate, the high labeled fraction in ^13^C-formate is possibly due to re-assimilation of ^13^CO2 by pyruvate carboxylase after formate oxidation by formate dehydrogenases (29). A similar jump in labeling in ^13^C-formate was observed for threonine and glutamate, which can both be derived from aspartate (Figure S11B,C). Alanine, on the other hand, had <5% labeling in ^13^C-formate like serine, consistent with also being derived from pyruvate (Figure S11B,C). Overall, no amino acid examined had labeling indicative of formate assimilation by the introduced pathway.

The analysis above requires a sufficiently high formate reduction flux to result in labeled proteins. For a more sensitive test of formate assimilation, we monitored fructose-6-phosphate (F6P), a metabolite immediately downstream of formaldehyde assimilation into the RuMP pathway. We added 200mM ^12^C- or ^13^C-formate to mid-exponential-phase cultures of control and pathway strains, continued incubating the cultures for 2 hours, and then harvested and extracted metabolites for LC-MS. We saw 8-10% labeling of F6P in ^12^C-formate, close to the expected background rate of 6% (Figure 5F, right panel). Labeling in ^13^C-formate was higher, at around 10% in all 3 strains. However, as in the case of the amino acids, there was no increase in the difference in labeling between labeled and unlabeled formate conditions (Figure 5G, right panel). Therefore, we were unable to detect evidence of our pathway assimilating formate into biomass *in vivo*.

It is possible that our ACS and ACDHs still do not have the activity needed to supply even detectable formaldehyde flux through the RuMP pathway. To test this, we used flux balance analysis (FBA) to calculate the theoretical growth rate that could be supported by the measured rate of formate reduction in the Nash assay. A genome-scale model of *M. flagellatus* KT metabolism does not exist, but we used a model developed for another RuMP-pathway methylotroph, *Methylotuvimicrobium buryatense* 5GB1C (30). We assumed that the flux through the methanol dehydrogenase reaction, which provides all the formaldehyde (and reduced carbon) for biomass production, is the same as the highest specific activity we measured in *M. flagellatus* KT lysates, or 1.2 nmol/min/mg (Figure 5D). We found that this would support a theoretical growth rate of 0.00048 h^-1^, or a doubling time of 8.6 weeks, even without ATP maintenance (with ATP maintenance, growth was infeasible). This is much slower than even the 55-hour growth supported by an unoptimized reductive glycine pathway (9), indicating that further improvements to activity and/or expression are needed.

## Discussion

### Utility of phylogenetically diverse enzymes

Previous work showed that EcACS and LmACDH have formate-reduction activity and that the enzymes are functional when expressed in *E. coli*. We extend that work to identify a panel of natural and engineered ACS and ACDH variants with improved expression and lysate activity and show that StACSstab6, MhACS2, and BmACDH, are well-expressed and active in the methylotroph *M. flagellatus* KT. None of these three enzymes had the highest ACS or ACDH activities *in vitro*, showing that expression in the host cytosolic environment is an equally if not more important factor than catalytic properties in practice. The computational design of StACSstab and the thermophilic source organisms of MhACS and BmACDH may have played a role in their greater expression and host range. By contrast, neither EcACS and LmACDH, the previous best enzymes for this pathway, were expressed in *M. flagellatus* KT, despite EcACS having the 2^nd^-highest formate *k_cat_*/*k_m_* of the wildtype ACSs and LmACDH2 being the most active ACDH we found. This highlights a key advantage of screening phylogenetic diversity in that this approach offers not only the chance to discover high activity, but also high expression and evolvability (20,31,32).

### Improving expression versus activity

Since we performed directed evolution on ACSs using a lysate-based screen, it is reassuring that we obtained increased lysate activity (5.8-fold for StACSstab and 1.9-fold for MhACS). However, this was entirely due to increases in functional expression (8-fold for StACSstab and 4.6-fold for MhACS) and not catalytic efficiency. In fact, *k_cat_*/*k_m_* decreased by 28% for StACSstab6 on formate, although it decreased by 72% on acetate, leading to an overall increase in the formate specificity from the parent enzyme. Despite this lack of improvement in catalytic activity, the increased soluble expression proved crucial to functionality in *M. flagellatus* KT, where StACSstab6, but not the StACSstab parent, was expressed and active. Interestingly, even wildtype ACS homologs differed widely in soluble expression in *E. coli* as well as in *M. flagellatus*. By contrast, there were no obvious differences in expression between the various ACDH homologs or evolved variants in *E. coli*, and our directed evolution of ACDH using a lysate-based assay led to increases in both lysate and specific activity.

Why did our lysate assays select for increased specific activity in ACDH but not in ACS? The ACS parents we chose perhaps started with poor stability or expression, but this is unlikely given their origins. Moreover, stability usually *decreases* while evolving for activity (33). A more likely possibility is that ACS activity is toxic. This has been observed previously (34), and thus decreasing it may allow cells to tolerate increased expression. This is consistent with the expression gain concomitant with a sudden reduction of acetate activity from StACSstab to StACSstab1/2 (N521V/L). Despite having a much lower acetate activity, however, even StACSstab6 still appears to be toxic, frequently leading to *E. coli* colonies with spontaneously decreased activity (one such colony can be seen as a replicate in Figure 3B). This problem can be mitigated in future rounds of evolution by using a low-background expression host and/or reducing induced expression level.

### Challenges of one-carbon substrates

A more fundamental problem is the possibility of biophysical limits on the formate activity of ACS. We chose ACS for synthesizing formyl-CoA because formate is structurally similar to ACS’s native substrate acetate. However, formate is less electrophilic than acetate, which could make it challenging to achieve a high *k_cat_*. Indeed, formate *k_cat_*s among our natural and evolved ACSs never exceeded 12 s^-1^, while the highest acetate *k_cat_* was 43.2 ±3.1 s^-1^. One-carbon compounds also have relatively few functional groups for interacting with a substrate-binding pocket, leading to higher *k_m_*s (35) and potentially explaining why even our lowest ACS *k_m_* for formate is greater than 40mM. As a result, our highest formate *k_cat_*/*k_m_* values are between 100-200 M^-1^s^-1^, 2-3 orders of magnitude lower than many natural enzymes, including the acetate activity for native ACSs. Encouragingly, however, natural enzymes find formate equally challenging. The *M. extorquens* formate-tetrahydrofolate ligase (FTL), which activates formate for assimilation, has a *k_m_* of 22 mM and a *k_cat_* of ∼100 s^-1^, for a *k_cat_*/*k_m_* of ∼5000 M^-1^s^-1^, about 30-fold higher than our best ACSs (36). Despite being 2 orders of magnitude lower than the median enzyme (35), this activity can support fully formatotrophic growth in natural and engineered pathways. Therefore, a physiologically relevant activity of the formate reduction module may be within reach given further enzyme engineering.

ACDH is not expected to be as challenging an engineering target as ACS, because most of the substrate binding affinity is contributed by the CoA group. We did not directly measure the *k_m_* of ACDH for formyl-CoA, but the ACS-coupled assays show it is at most 7 mM (Figure S7). In reality it is probably much lower; the *k_m_* of ACDH for acetyl-CoA can be <100µM (37), and the *k_m_* of 2-hydroxyacyl-CoA lyase for formyl-CoA is 200µM (15), despite this not being its native substrate. However, a potential problem with ACDH is *k_cat_*. Even though the fastest ACDH homolog in the literature has a *k_cat_* of ∼60 s^-1^ on acetate (38), our best ACSs were almost 2 orders of magnitude slower on formate. However, given that we were able to increase this value by ∼2-fold in 2 rounds of directed evolution, further engineering will likely result in additional gains.

### Effects of mutations on ACS and ACDH

Previous work found that ACS substrate specificity can be changed from acetate to larger or more polar substrates by mutating V310, T311, V386, or W414 in StACS, which are all with 4Å of the acetyl moiety (20,22,23). However, for formate, we did not find increases in lysate activity when mutating V310, T311, or V386 in isolation or V310 and V386 combinatorially (Table 1).

Instead, we found that the previously unexplored N521V/L mutations cause a large decrease in acetate activity in StACSstab. In the 2p2f structure of StACS, acetate is modeled in the active site with its acetyl carbon pointed toward N521, 5.7 Å away from the asparagine’s sidechain carbonyl (23). This orientation is consistent with the ability of large hydrophobic substitutions at N521 to favor a smaller acyl moiety in the substrate.

Several other mutations increased ACS lysate activity while decreasing acetate *k_cat_*/*k_m_*. StACSstab F260W and W414F increased acetate *k_m_*, as can be seen by comparing StACSstab3 and StACSstab4 to StACSstab1. In the StACS structure, W414 is within 4 Å of the acyl substrate, while F260 is 10.9 Å away but in contact with the first-shell V310 sidechain (Figure 3A). MhACS Y499V decreased acetate *k_cat_*; it is 11.1 Å from the acyl substrate but contacts the backbone of W414. The homologous StACSstab Y496 had mutants among the top hits in the first round of directed evolution (Figure S5B), but was not pursued in favor of N521V. StACSstab G524 lines the CoA binding site in StACSstab and as a result, G524S/L is known to block CoA addition to the acyl group(23). Our results show that G524A, on top of N521V W414F, maintains formate activity while increasing acetate *k_m_*. Additional residues (e.g. F421) showed evidence of improved activity in our initial screens on StACSstab (Figure S5), but were not pursued fully. They are prime candidates for mutagenesis in future rounds of evolution.

Over two rounds of site-saturating mutagenesis and screening at 9 residues comprising the acyl-binding tunnel in LmACDH, we found a double-mutant LmACDH2 (A252S S253C) that contributed to improved activity on formyl-CoA. These positions partially overlap with those mutagenized in a recent effort to engineer an ACDH to reduce glycolyl-CoA to glycolate (20). Our evolved isolates were comparable in activity to the best natural homologs BmACDH and PtACDH, but ultimately only BmACDH expressed well in *M. flagellatus* KT. BmACDH has the same sequence as LmACDH at positions homologous to A252 and S253, so these are clear candidates for mutagenesis in future directed evolution efforts.

### *M. flagellatus* KT as an *in vivo* pathway testing platform

We chose to use *M. flagellatus* KT for testing *in vivo* activity of our pathway because it could potentially gain formatotrophy with only the expression of ACS and ACDH. This is the first published instance, to our knowledge, of metabolic engineering in this organism. Recently, an *E. coli* strain was engineered to grow on methanol as a sole carbon source via the RuMP pathway (39). This provides the alternative option of engineering the ACS/ACDH pathway in *E. coli* instead, which would allow access to a wider range of genetic tools and pathway manipulations (40). Most importantly, it would allow the improvements we obtain from directed evolution in *E. coli* lysates to be directed translated into *in vivo* activity or expression. However, even in the fully methylotrophic *E. coli*, formaldehyde toxicity is still a major problem, and perhaps as a result, its doubling time on methanol is more than 8 hours. By contrast, *M. flagellatus* KT has a doubling time of 2 hours on methanol, indicating its naturally evolved robustness against formaldehyde toxicity. Perhaps the best approach in future work is to use a combination of strains – *E. coli* for initial troubleshooting and improvement of enzymes, followed by *M. flagellatus* KT for fine-tuning for maximal flux.

## Conclusion

Through phylogenetic homolog screening and directed evolution, we identified a panel of highly expressed and active ACS and ACDH enzymes and gained insight into the genetic determinants of their acyl-substrate specificity. We established a plasmid-based expression system in the RuMP-pathway methylotroph *M. flagellatus* KT and used it to introduce a formate-reduction pathway in an attempt to confer synthetic formatotrophy. Although we ultimately did not observe *in vivo* formate assimilation via our pathway, the enzymes and insights from this work should enable continued improvement of this pathway toward the ultimate goal of efficient conversion of CO2 into value-added chemicals.

## Materials and Methods

### Bioinformatics and enzyme homolog selection

All bioinformatics and analysis/visualization of experimental data was performed in Python/Jupyter. Phylogenetic trees for figures were computed by FastTree (41) and visualized using iTOL (42).

To identify ACS homologs for testing, an initial candidate list of 8,911 sequences was compiled that included: 6,104 sequences from the “Acetate-CoA ligase” Interpro family (IPR011904)(43) with the same 3 domains as EcACS downclustered to 90% identity using CD-HIT (44); 2,790 sequences from RefProt based on a pHMMER search (E-value < 10^-200^) with query EcACS (P27550) (45); 17 experimentally characterized ACS homologs from BRENDA (EC 6.2.1.1) (46). From the initial candidate list, an alignment and distance matrix was generated using Clustal Omega (47), and a hierarchical clustering of the distance matrix was used to guide manual selection of a diverse final set of ACSs. Initially 11 ACSs were chosen for testing (see below for details). Then, given the results of the 1^st^ round of testing, 30 additional ACSs were chosen to further sample clades with high activity while also exploring new areas of sequence space.

For ACDHs, 7,646 sequences were obtained from UniProt via a pHMMER search (E-value < 7×10^-17^) with query LmACDH (Q8Y7U1) or enzyme commission number search (EC 1.2.1.10), or from BRENDA (EC 1.2.1.10). After alignment and clustering of ACDHs, mhpF-like sequences were removed, leaving 4,037 adhE-like sequences from which the final selection was made.

Initially 9 ACDHs were chosen for testing; then these results were used to choose 37 additional ACDHs to test. The full phylogeny of all ACSs and ACDHs considered for homolog discovery would be too large to visualize as a tree, so Figure S1 shows only untested sequences that have less than 50% (ACSs) or 40% (ACDHs) amino-acid identity to each other and to the tested homologs. Some of the tested homologs are more similar to each other than this because they were chosen for reasons other than diversity; these were all included in the trees.

### DNA synthesis and *E. coli* strain construction

ACS and ACDH sequences were codon optimized for *E. coli* expression using Integrated DNA Technologies’ online tool (https://www.idtdna.com/CodonOpt; accessed September 2018). DNA synthesis was performed at Twist or the Joint Genome Institute of the U.S. Department of Energy. Genes were cloned into vector pET29b+ between *Nde*I and *Xho*I such that expressed enzymes have a C-terminal 6xHis tag. Expression vectors with ACDH genes were electroporated into *E. coli* expression strain BL21*(DE3), propagated on lysogeny broth (LB) + 50ug/mL kanamycin, and stored at −80C in 25% glycerol.

ACS is known to be repressed under standard physiological conditions by acetylation at K609, but can be derepressed by a point mutation L641P (48). We found that a simpler method of knocking out the patZ deacetylase leads to comparable EcACS activity (Figure S2A), so we used a BL21*(DE3) ΔpatZ host strain for all ACS experiments. The *patZ* gene was deleted from BL21*(DE3) using lambdaRed recombinase (49). For the round of directed evolution from MhACS2 to MhACS3, the alternate expression strain NovaBlue(DE3) ΔpatZ was constructed and used.

### Protein expression, lysis, and SDS-PAGE

To express proteins, strains were inoculated directly from −80C stocks into auto-induction medium (50) at 1:500 to 1:50,000 dilution and incubated at 37C for 24 hours. For screening, 500µL cultures were grown in 2mL 96-well microtiter plates (Axygen P-DW-20-C) with shaking at 1000rpm on a benchtop shaker (Heidolph Titramax 1000) in a temperature-controlled room.

For SDS-PAGE and Nash assays, 5mL cultures were grown in round-bottom glass tubes in a rotary shaker incubator at 250rpm. For purification, 50mL or 500mL cultures were grown in Erlenmeyer flasks and shaken at 250rpm.

To prepare lysates for screening, 96-well plate cultures were pelleted at 2200g for 10 min, washed once in 4°C water, and resuspended by vortexing after adding 300µL/well of lysis buffer (50mM HEPES pH 7.5, 50mM NaCl, 2 mM MgCl2) with 0.6 mg/ml lysozyme, 0.1mg/mL polymyxin B, and 1:50,000 Sigma benzonase nuclease. Plates were incubated at 37°C for 50 min without shaking followed by 10 min shaking at 1000rpm. Then, lysates were pelleted at 2200g for 10 min, and the supernatant was used for downstream assays.

To prepare lysates for SDS-PAGE, Nash assay, or protein purification, 5mL, 50mL, or 500mL cultures were pelleted at 4000rpm for 10min and resuspended, respectively, in 0.5mL, 8mL, or 30mL lysis buffer with 0.1mM DTT, 1:500 Sigma protease inhibitor cocktail, 1:50,000 Sigma benzonase nuclease. Cell suspensions were sonicated on ice (Branson SLPt) for 3 repeats of 10 seconds on and 10 seconds off at 30% amplitude for 5mL cultures, or 6 repeats of 30 seconds on and off at 70% amplitude for 50mL cultures, or 12 repeats of 30 seconds on and off at 70% amplitude for 500mL cultures. Lysates were pelleted at 4000rpm for 15 minutes and the supernatant used for analysis or purification.

To analyze lysates and purified enzymes by SDS-PAGE, samples were mixed 1:1 with 2x Laemmli sample buffer with 2-mercaptoethanol (Bio-Rad) and boiled for 10min. A sample containing 1-20µg of protein was loaded into a 4-15% precast gel (Bio-Rad Mini-ProTEAN) and run at 60V for 20min followed by 160V for 1 hour. To compare expression levels across lysates, protein concentration was determined by BCA and equal µg of protein were loaded in each lane. Gels were stained by Coomassie blue and imaged using a digital camera. Minor contrast adjustments were made to the images to improve visibility of bands.

### Assays for enzyme activity in lysates

ACS was assayed in lysates using a discontinuous assay with DTNB (5,5-dithio-bis-(2-nitrobenzoic acid)), which reacts with CoA to yield absorbance at 412nm (20, 51). In a microtiter plate (Corning Costar 3370), 100µL of reaction buffer (10 µL of expression-induced *E. coli* lysate, 50mM HEPES pH 7.5, 2mM MgCl2, 5mM ATP, 0.5mM CoA, and 50mM sodium formate) was aliquoted. The formate was added last to start the reaction, everything was incubated for 10min at 37°C, then stopped by adding 100µL DTNB reagent (50mM HEPES pH 7.5, 2mM DTNB). Absorbance at 412nm was read on a plate reader (Molecular Devices Spectramax 190). Empty vector control lysates were used to establish the background signal, and the metric *ΔA412 = A412_Empty Vector_− A412* was used to quantify lysate ACS activity. Note that higher ACS activity corresponds to lower *A412* but higher *ΔA412*. For the first round of ACS directed evolution, a continuous assay was used (see “Protein purification and enzyme kinetics” below), but subsequent rounds of evolution used the discontinuous assay described above.

ACDH lysates were assayed in a continuous assay by coupling to ACS, all steps at 37°C. Reactions were performed in a microtiter plate with a total volume 200µL containing 2µL clarified lysate, 2µM StACSstab1, 50mM HEPES pH 7.5, 5mM MgCl_2_, 1mM DTT, 2.5mM ATP, 0.5mM CoA, 0.6mM NADH, and 50mM formate. Reactions were prepared in 100µL at 2x concentration and then 100µL of 2x formate was added to start the reaction. Absorbance at 340nm was monitored and NADH concentration was calculated as [*NADH*] = ^A340/ε.l^ where *ε = 6.22 mM^-1^ cm^-1^* is the extinction coefficient of NADH and *l* = 0.56 *cm* is the path length of 200µL of reaction mixture in the microtiter plate. Initial velocities were calculated from least-square linear fits to the first 3-10 datapoints. The amount of ACS to use for coupling was determined by titrating ACS for every new batch of purified ACS or round of lysate screening (Figure S5). For assaying purified ACDHs, we used 30x molar excess of coupling ACS.

### Directed evolution to improve ACS and ACDH

To engineer StACSstab and MhACS, residues were selected for site-saturating mutagenesis based on proximity to the acetate molecule in the crystal structure of the *Salmonella typhimurium* ACS (PDB: 2p2f). Mutant libraries at single positions were constructed using “inside-out” PCR from NNK or “22c” (52) degenerate primers and multi-site combinatorial libraries were made by overlap-extension PCR. Libraries were electroporated into expression host strains (see above). One 96-well plate of mutant clones (plus control strains) was screened for each single-site library. Eight plates were screened for the 4-site library in MhACS evolution round 1. The best 10-20 mutants were restreaked on LB + kan plates and 4 colonies of each mutant were screened again. The best mutant from the secondary screen was used as the parent for the next round of evolution. The DTNB assay with 50mM formate was used for all screening all ACS mutants, except in round 1 of StACSstab evolution, when the myokinase coupled assay was used. To engineer LmACDH, its crystal structure (3k9d) was superimposed on the *R. palustris* ACDH (5jfn) (53), and the position of the substrate propionyl-CoA from 5jfn was used to choose residues in 3k9d for saturation mutagenisis. Screening was done using the ACS-coupled assay described above on clarified lysates at 50mM formate.

### Protein purification and enzyme kinetics

All steps were done at 4°C. 1mL of Ni-NTA superflow resin (Qiagen) was placed in a gravity-flow column (GE Healthcare PD-10) and equilibrated by flowing through 10mL of lysis buffer. Then 8mL of clarified lysate was added and the column was sealed and placed on ice and nutated (VWR 12620-916) for 10 minutes. Then the lysate was flowed through the column, 10mL of wash buffer (50mL HEPES pH 7.5, 300mM NaCl, 35mM imidazole) was applied, and protein was eluted in 10mL of elution buffer (50mM HEPES pH 7.5, 50mM NaCl, 150mM imidazole). Eluate was exchanged to lysis buffer by spinning at 4000g for 15min in Amicon Ultra-15 30kDa (for ACS) or 10kDa (for ACDH) centrifugal filters. Glycerol was added to 10% and protein concentration was determined by BCA assay (Pierce 23227). Kinetic assays were performed immediately after purification. Additional purified enzyme was split into aliquots and stored at −20°C.

For the first round of ACS directed evolution, a secondary screen was performed with high-throughput purification. 1mL cultures of *E. coli* expression strains were lysed in 300µL lysis buffer + 1:50,000 benzonase + 0.6mg/mL lysozyme + 0.1mg/mL polymyxin B and clarified lysates were flowed over 50µL Ni-NTA superflow resin in each well of a 96-well filter plate (Pall) and by centrifugation at 2200g for 10 min. The resin was washed with 200µL of wash buffer and eluted in 100µL elution buffer. Eluates were used immediately without buffer exchange and protein was quantified by BCA.

ACS kinetics were determined by a continuous assay using coupling enzymes (23, 51). All steps were performed at 37°C. All coupling enzymes were from Sigma. A reaction buffer was prepared with 0.05 – 0.2µM of ACS, 15 U/mL pyruvate kinase, 23 U/mL lactate dehydrogenase, and 25 U/mL myokinase in 50mM HEPES pH 7.5, 5mM MgCl_2_, 1mM DTT, 0.6mM NADH, 2.5mM phosphoenolpyruvate, 2.5mM ATP, and 0.5mM CoA. 100µL of a 2x portion of the reaction mixture was aliquoted into a microtiter plate and the reaction started by adding 100µL of 2x sodium formate or acetate. Absorbance at 340nm was monitored for 10 minutes on the plate reader and initial velocities of NADH oxidation were calculated as above. Kinetic parameters *k*_cat_ and *k*_m_ were extracted from plots of initial velocities versus substrate concentration by fitting, where *v* is the per-enzyme initial velocity, [*S*] is the substrate concentration, and *b* is a background rate. Kinetic curves were fit using `scipy.optimize.curve_fit`. Purified ACDHs were assayed similarly to ACDH lysates as described above, except rates of NADH oxidation were normalized to enzyme concentration as determined by BCA.

### Cloning and *M. flagellatus* KT strain construction

All genetic constructs for *M. flagellatus* KT expression were maintained on a IncP-based broad-host-range plasmid, whose backbone was derived from pAWP87 / pCM66 (27). Expression vectors for *M. flagellatus* KT were constructed using PCR amplified backbone fragments, promoters from the *M. flagellatus* KT genome, and ACS/ACDH coding regions from DNA synthesis. PCR primers were designed with 20-25bp of overlap and Gibson assembled (New England Biolabs) and electroporated into *E. coli* strain TOP10. Electrotransformants were propagated in LB + 50ug/mL kanamycin at 20°C to avoid toxicity of the constructs.

A rifamycin-resistant isolate of *M. flagellatus* KT was used for triparental conjugation. Briefly, *M. flagellatus* KT was patched, along with plasmid donor strain and helper strain pRK2073 (54), onto plates containing MM2 medium (14.5 mM K_2_HPO_4_, 18.8 mM NaH_2_PO_4_ (monohydrous), 0.8 mM MgSO_4_ (heptahydrous), 3.8mM Na_2_SO_4_, 9.9mM KNO_3_, and 1x Vishniac trace elements (55)) with 5% LB and 2% methanol and incubated 16-20 hours at 37°C. Cell mixtures were then plated on plates with MM2 + 0.1% pyruvate + 2% methanol + 50ug/mL rifamycin + 50ug/mL kanamycin and incubated for 2-3 days at 37°C, colonies were restreaked onto the same medium and incubated for another 2-3 days at 37°C. For each conjugation, multiple colonies were screened and by SDS-PAGE and Nash assay and the colony with the best phenotype was used for downstream assays. Strains were stored by culturing in liquid MM2 medium and then freezing at −80°C with 10% DMSO.

Promoter reporter constructs were constructed by Gibson assembly of various promoters upstream of dTomato in pAWP87. These were conjugated into *M. flagellatus* KT, colonies were inoculated into seed cultures in MM2 + 2% methanol + kanamycin, grown for 24-48 hours at 37°C with shaking, diluted 1:50 and grown 24 hours, and then measured at 535nm excitation / 590nm emission on a plate reader (Tecan Infinite 500). Four promoters (Phps, PmxaF, Ptrc, Ptac) were chosen for driving ACS/ACDH, but intact plasmids with Ptrc and Ptac could not be isolated in the *E. coli* TOP10 cloning strain and were omitted from further experiments.

### Nash assay

*M. flagellatus* KT strains were patched from −80°C stocks onto MM2 + 2% methanol + 50ug/mL kanamycin (“MM2 Me2 kan”) plates and incubated at 37°C for 2 days, then inoculated into 5mL liquid MM2 Me2 kan medium in a round-bottom glass tube and incubated at 37°C with 250rpm shaking for 24 hours. Cultures were lysed as described above and the soluble fraction was used for the assay. In a microtiter plate, 8µL lysate was added to a reaction mixture (50mM HEPES pH 7.5, 5mM MgCl_2_, 1mM DTT, 4mM ATP, 6mM NADH, and 1mM CoA, either 4µM ACDH, 2µM ACS, or no additional enzyme, and either 0mM or 300mM formate) for 100µL total volume. Reactions were incubated for 1 hour at 37°C. Then 100µL of Nash reagent (0.1M ammonium acetate, 0.2% acetic acid, 3.89M acetylacetone) is added and reactions are incubated at 65°C for 30 min. Then, to precipitate proteins, 20µL of 100% w/v trichloroacetic acid is added and the reactions are placed on ice for 5 min. The plate is spun down at 2200g for 10min and 100µL of supernatant is transferred to a new plate and *A412*nm is measured. The difference between absorbance at 300mM and 0mM formate, *ΔA412* = A412_300mM_− A412_0mM_, was used to quantify lysate activity.

### ^13^C formate labeling and analysis by LC-MS

To analyze proteinogenic amino acids, *M. flagellatus* KT strains were revived from −80°C and grown in 5mL liquid MM2 Me2 kan seed cultures as described above. 200µL of seed culture was transferred to 5mL of MM2 + kan + 0.05% methanol + 200mM ^12^C or ^13^C formate. Initial experiments used cultures with one growth cycle in MM2 + 2% methanol + 200mM formate. Later experiments used multiple growth cycles as follows. Cultures were grown for 24 hours, pelleted at 2200g for 10min, and resuspended in fresh medium with methanol and formate. This iterative re-feeding was done for 3 days, and the final cell pellets were resuspended in 1mL 6N hydrochloric acid and boiled in glass vials for 24 hours (56). The vials were uncapped and left to dry for another 24 hours. The biomass was then resuspended in 1mL of water, dried for 24 hours, and then resuspended in 0.5mL of water and centrifuge-filtered (Costar Spin-X 0.22 μm, Sigma). Samples were stored at −20°C until LC-MS.

LC-MS was performed as in (57). A Waters Xevo mass spectrometry triple quad (Xevo, Waters, Milford, MA) with UPLC system equipped with a Zic-pHILIC column (SeQuant, PEEK 150 mm length × 2.1 mm metal free, with 5 μm polymeric film thickness, EMD Millipore) was used for detection of MID of metabolites with following LC condition. Mobile phase A is 20 mM bicarbonate in water (OPTIMA grade, Thermo Fisher Scientific), mobile phase B is 100% acetonitrile (OPTIMA grade, Thermo Fisher Scientific). The LC condition starts with 0.15 ml/min flow rate with initial gradient A = 15% for 0.5 min, the increased to 80% A in 20 min, at 21 min, A = 90%, and hold for 5 min, at 26.5 min, mobile phase A switched to 15% and then re-equilibrate the column for 5.5 min. Multiple reaction monitors (MRM) were set up for each metabolite of interest. For each metabolite, 12C chemical standards were used to set up the mass channel for unlabeled isotopomers. The predicted mass fragments were then used to predict the MRM for labeled isotopomers for each metabolite. The MassLynx software (Waters) was used to integrate ion peak intensities, with subsequent analysis in Python.

### Availability of Materials

All strains and plasmids used in this study are available upon request from the authors, and all plasmids other than the homolog screening libraries have been deposited in Addgene.

## Acknowledgements

We thank Yakov Kipnis, Devin Trudeau, Joseph Groom, Julius Palme, and Dan Tawfik for valuable technical advice and feedback on the manuscript.

## Data availability

All homolog protein sequences, plasmid sequences, and processed data on lysate and purified enzyme activities are contained in the supplementary tables. Raw data on enzyme activities and Python code used for analysis are available from authors upon request.

## Funding

This work was supported by a Washington Research Foundation Postdoctoral Fellowship to J.W. and funding from the University of Washington to M.E.L. Gene synthesis was provided by the Synthetic Biology program at the Joint Genome Institute of the Department of Energy.

## Conflicts of interest

The authors declare no conflicts of interest.

## Notes

### Competing Interest Statement

The authors have declared no competing interest.

### Summary of Updates

The author list was corrected. No other changes have been made.

