## Supplemental Information for "Enzyme engineering and *in vivo* testing of a formate-reduction pathway"

**Table S1. Plasmids and strains used in this study**

See Excel file.

**Table S2. ACS and ACDH homolog sequences**

See Excel file.

**Table S3. ACS and ACDH homolog screening data**

See Excel file. A) Raw A412 values from the endpoint assay are provided. All enzymes are screened in BL21*(DE3) ΔpatZ E. coli expression strain unless denoted "patZ+", which is BL21*(DE3). "Adjusted p-value" is a t-test p-value for difference from Empty vector adjusted for multiple testing across the homologs by the Benjamini-Hochberg procedure. B) NADH oxidation rates from ACS-coupled lysate assay are provided, in units of $\mu$M/s. "Adjusted p-value" is calculated as described above, using combined data from both isolates.

**Table S4. Plasmids and strains used in this study**

See Excel file. A) Means and standard deviations of kcat, Km, and kcat/Km values of the 6 ACS homologs chosen in Figure 1 and plotted in Figure S5. B) Means and standard deviations of kcat, Km, and kcat/Km values parent and evolved ACSs as plotted in Figure 2. This was a separate batch of purified proteins and kinetic assays from those in Table S2A, so enzymes in both datasets have slightly different values. C) NADH oxidation rates from ACS-coupled assay on purified ACDHs, in units of $s^{-1}$.

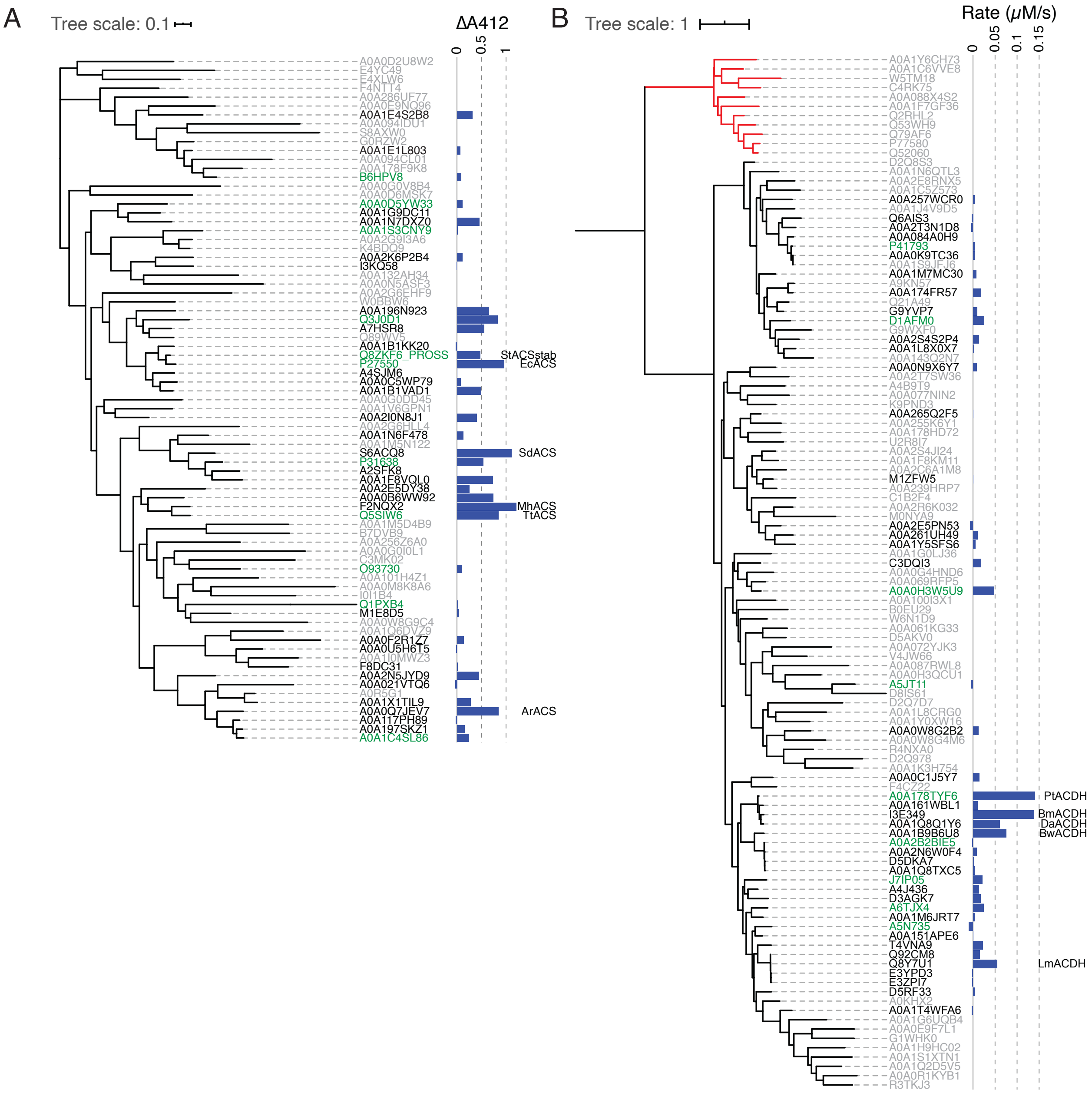


**Figure S1. Phylogenetic tree of ACS and ACDH homologs and activity screening results**

A) Phylogeny of 78 ACSs labeled by Uniprot accessions. The full set of 8,911 ACSs was downclustered to 50% amino-acid identity and combined with the assayed ACSs (Materials and Methods). 11 were assayed in the 1st round of homolog discovery (green font) and 30 in the 2nd round (black font), and 47 were not tested (gray font). ΔA412 bars on the right represent a measure of activity in clarified lysates with formate as substrate. B) Phylogeny of 111 ACDHs, generated and labeled as in (A), except sequences were downclustered at 40% identity to limit the size of the tree. 9 enzymes were tested in the

1st round, 37 in the 2nd round, and 65 were not tested. Clade in red represents aldolase-ACDH fusions (such as *E. coli* dmpF) that were not considered for testing. Activities in clarified lysates are shown on the right.

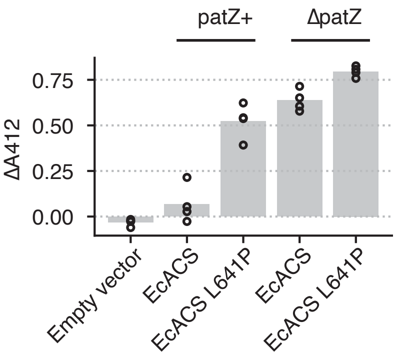


**Figure S2. Effect of ΔpatZ on ACS activity**

A) Lysate activity for wildtype *E. coli* ACS and the L641P mutant in wildtype BL21*(DE3) ("patZ+") and BL21*(DE3) ΔpatZ ("ΔpatZ") expression host strains from the discontinuous DTNB-based assay. Each bar is the mean of 3 replicates.

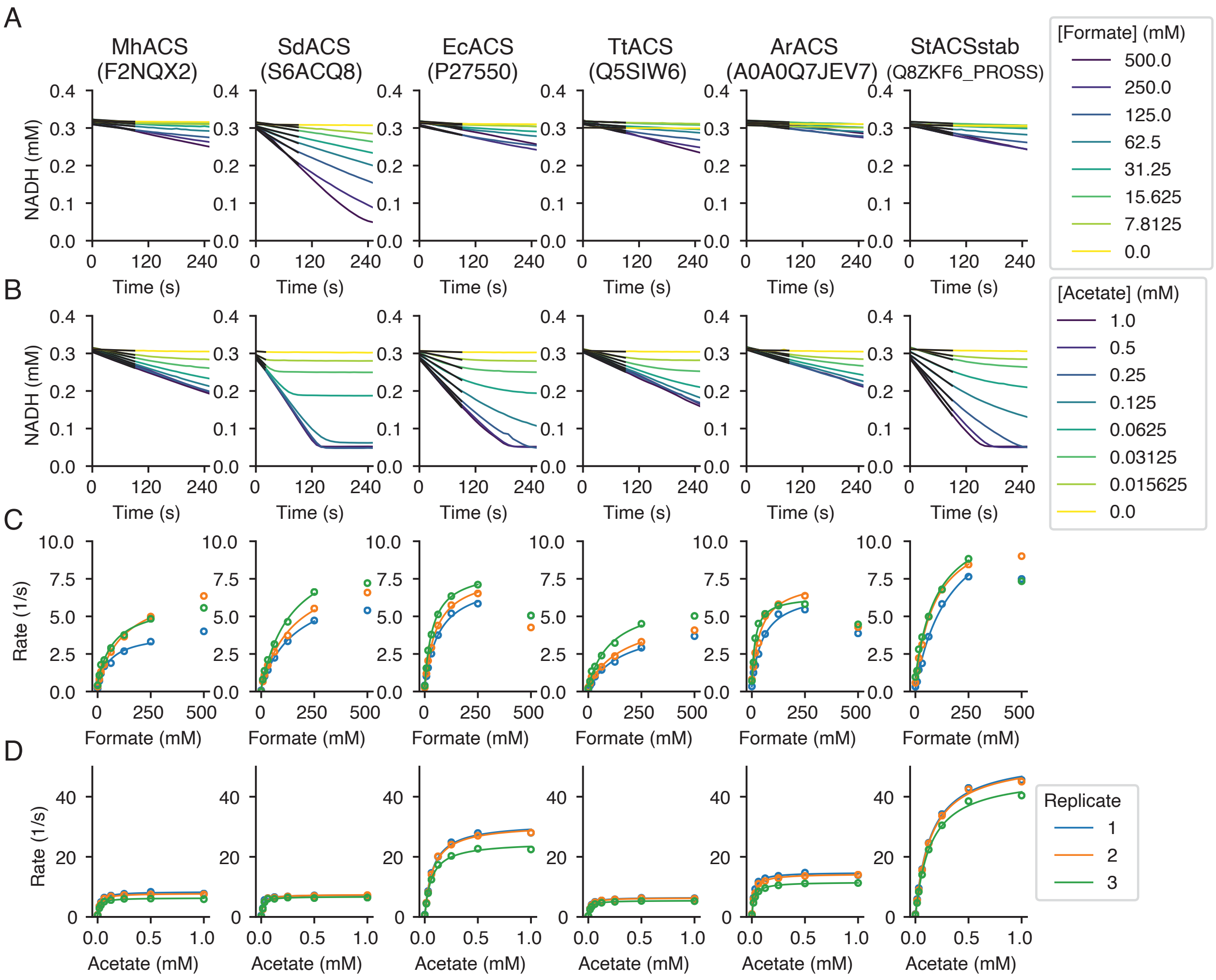


**Figure S3. Enzyme kinetics of selected ACS homologs.**

NADH versus time traces with formate (A) or acetate (B) as the substrate for the 6 ACS homologs chosen in Figure 1 for follow-up analysis, using an NADH-based continuous assay. A least-squares linear fit to the first 10 datapoints (black line; 3 points were used for SdACS in acetate) was used to extract the initial velocity, which is normalized to protein concentration and then plotted against formate (C) or acetate (D) concentration. A Michaelis-Menten curve with background was fitted to velocity versus substrate concentration, omitting the 500mM formate data point due to apparent substrate inhibition in some ACSs. These curve fits were used to determine the $k_{cat}$ and $K_m$ values plotted in Figures 2 and S4 and a similar procedure was used for Figure 3.

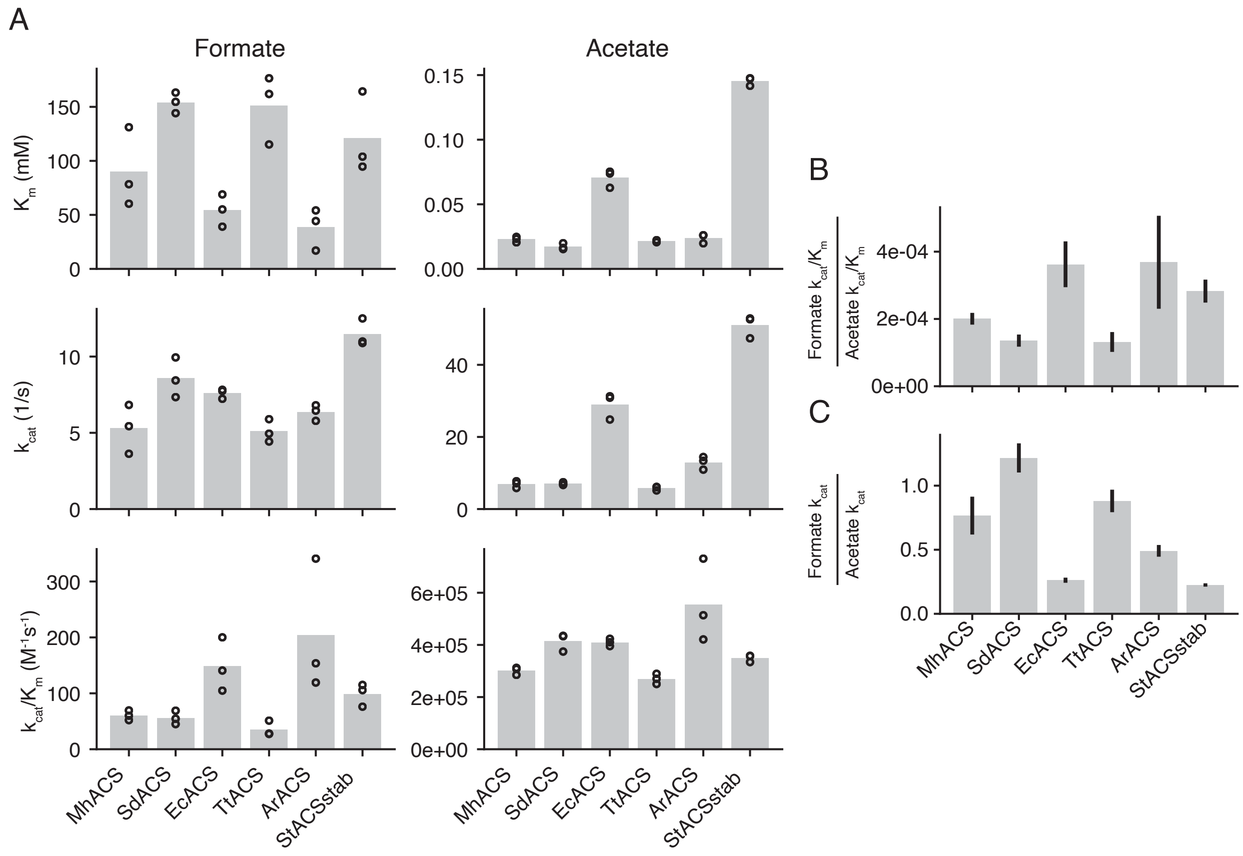


**Figure S4. Kinetic parameters of selected ACS homologs on formate and acetate**

A) $K_m$, $k_{cat}$, and $k_{cat}/K_m$ for formate and acetate of the 6 selected ACS homologs, extracted from the kinetic data shown in Figure S2. Circles show 3 replicates and bars show the mean. B) Ratios of formate $k_{cat}/K_m$ to acetate $k_{cat}/K_m$. C) Ratios of formate $k_{cat}$ to acetate $k_{cat}$. Error bars represent s.d. of the ratios estimated using the replicate data (see Materials and Methods).

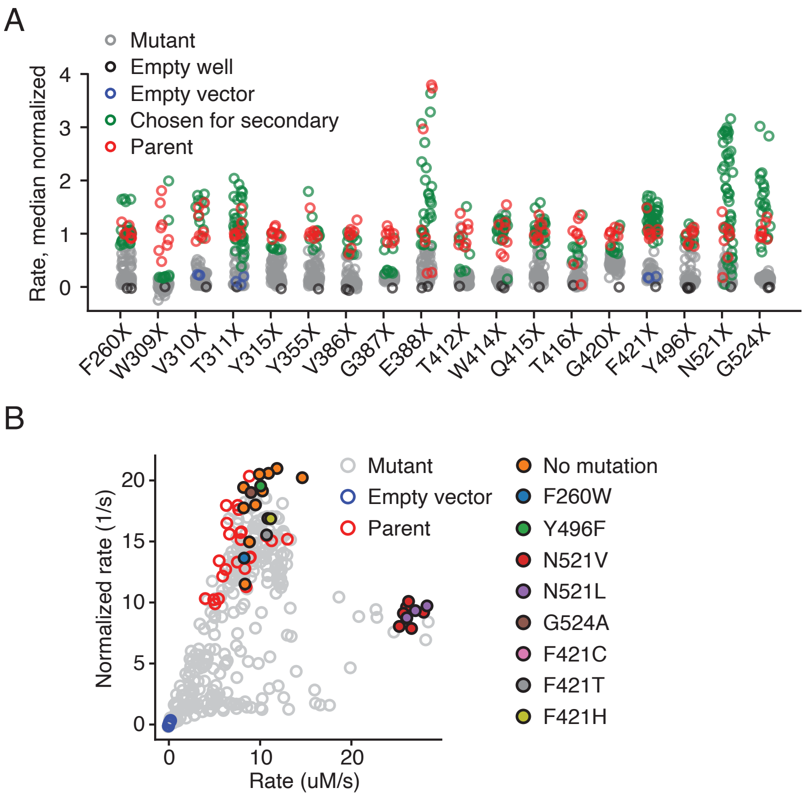


**Figure S5. StACSstab directed evolution round 1 screening**

A) Reaction rates of StACSstab site-saturating mutant libraries by the continuous assay in 50mM formate. The most-active mutants in each library were chosen for a secondary screen. B) Enzyme-concentration-normalized rates versus unnormalized rate in secondary screen with 50mM formate. The same assay was used as in (A) but enzymes were purified (Materials and Methods). Mutants with the highest total and normalized rates were Sanger-sequenced and labeled in colored circles.

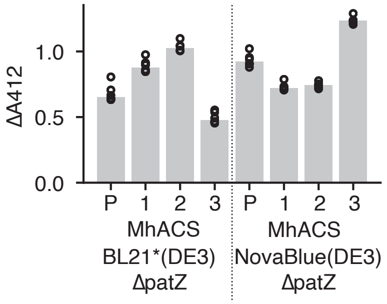


**Figure S6. Activity of evolved MhACS enzymes in different expression hosts**

Activity was assessed with the discontinuous assay. Circles show 3 replicates and bars show the mean. "P" indicates parental or wildtype enzyme, and numbered variants correspond to mutants listed in Table 1.

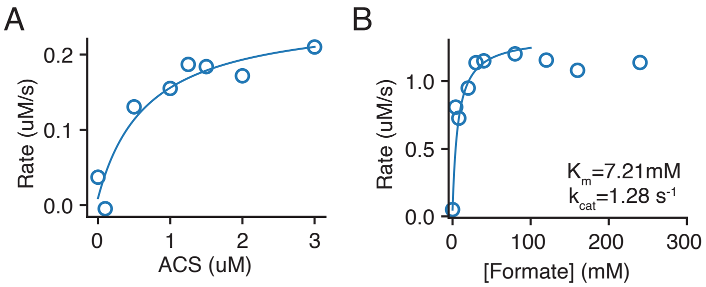


**Figure S7. ACS-coupled assay for ACDH activity**

A) Rate of NADH oxidation by LmACDH in the presence of saturating (350mM) formate, 2uL *E. coli* lysate, and various amounts of ACS. Curve is a least-squares Michaelis-Menten function. A titration of coupling ACS was performed for every new batch of purified ACS or round of lysate screening. For assaying purified ACDHs, we used 30x molar excess of coupling ACS. B) Rate of NADH oxidation by LmACDH versus formate concentration, with 30x coupling ACS. This shows that the $K_m$ of LmACDH is at most 7.2 mM.

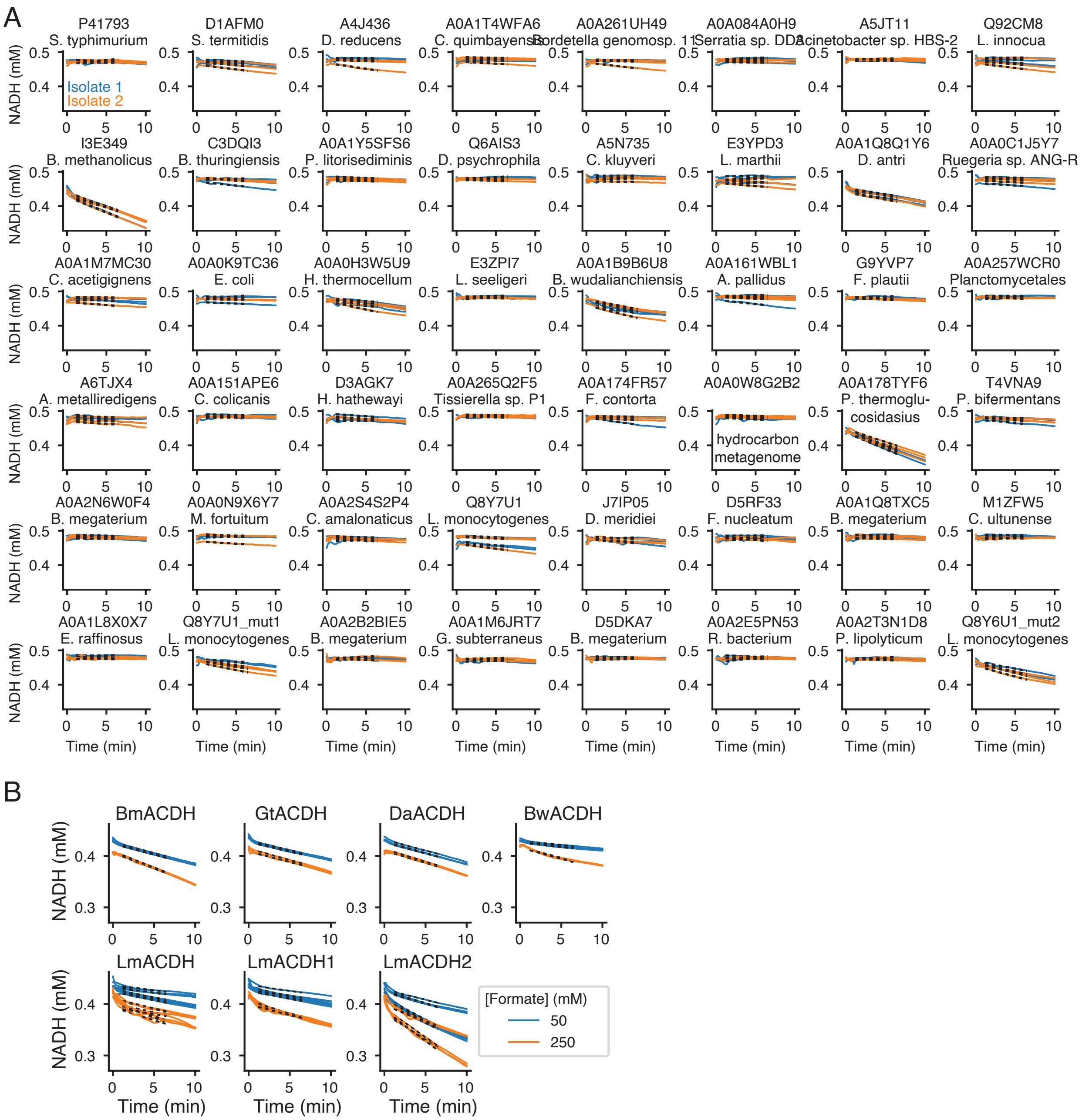


**Figure S8. NADH oxidation curves from ACDH activity measurements**

A) NADH oxidation curves of ACS-coupled assays of *E. coli* lysates expressing 46 ACDH homologs and 2 evolved mutants of LmACDH. Three replicates of 2 independent isolates are shown. Black dotted lines are least-squares linear fits to extract reaction rates, plotted in Figure 4A, C. B) NADH oxidations curves of ACS-coupled assays on purified ACDH homologs and evolved variants. Rates were extracted by linear fit and plotted in Figure 4D.

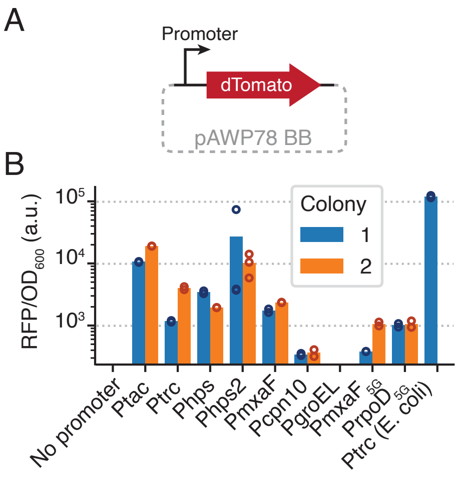


**Figure S9. Promoter validation for *M. flagellatus* KT**

A) Schematic of fluorescent protein reporter plasmid for testing promoters in *M. flagellatus* KT. B) OD-normalized fluorescence for a panel of promoters transformed into *M. flagellatus* KT strains. Mean (bars) and 3 replicates (circles) are shown for 2 independent transformant colonies. Subscript “5G” indicates promoter sequences from *M. buryatense* 5GB1. “Ptrc (E. coli)” is an *E. coli* strain containing the same plasmid.

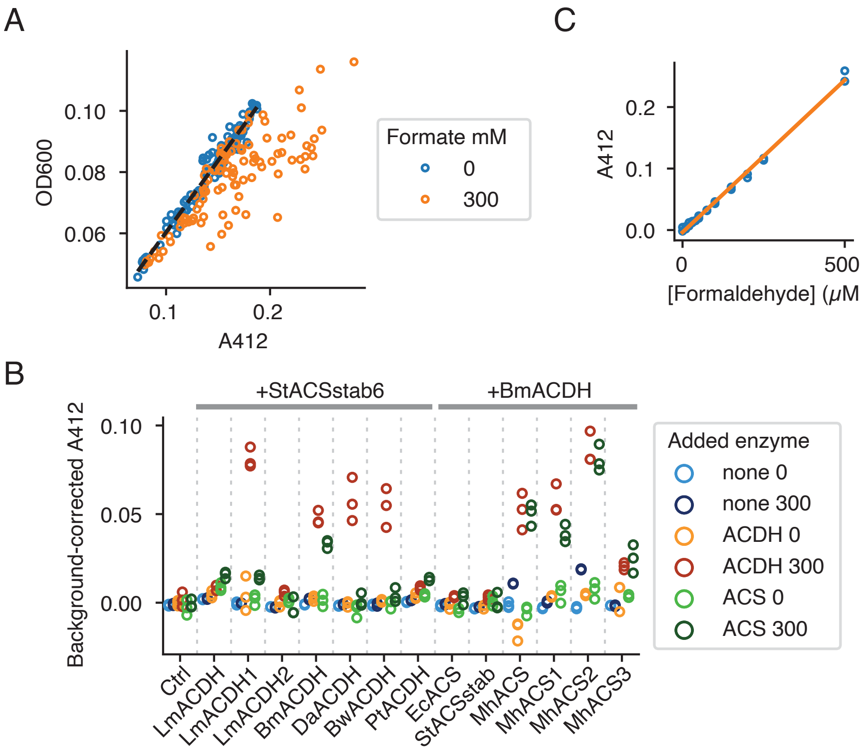


**Figure S10. Nash assay measurement of pathway specific activity**

A) Absorbance at 600nm (OD600) versus absorbance at 412nm (A412) for Nash assay samples after trichloroacetic acid precipitation. A least-squares linear fit to the data with no formate (and hence no expected Nash reaction) allows the background of each sample to be estimated and then subtracted from its measured A412. B) Background-corrected A412 of Nash assays on *M. flagellatus* KT lysates. Differences between 300mM and 0mM conditions were then converted to specific activities as shown in Figure 5D,E. C) Standard curve of A412 versus formaldehyde concentration used to convert A412 values into formaldehyde concentration for calculation of specific activity.

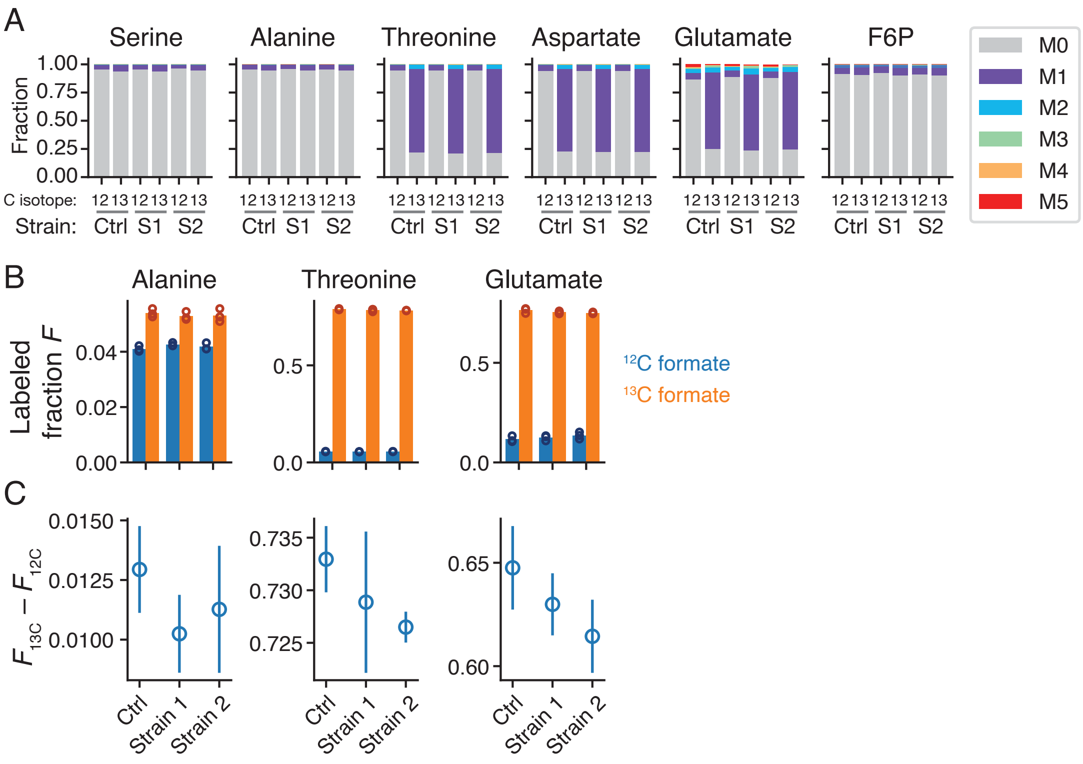


**Figure S11. 13C labeling of selected amino acids and fructose-6-phosphate**

A) Fraction of each metabolite labeled at 0 to 5 carbons ("M0" through "M5") for control strain, Strain 1, and Strain 2 grown in $^{12}$C- or $^{13}$C-formate. "Labeled fraction" in Figure 5F is simply 1 – (M0 fraction). B) Fraction of $^{13}$C-labeled proteinogenic alanine, threonine, and glutamate from cells grown in methanol + 200mM $^{12}$C or $^{13}$C formate. 3 biological replicates of the control strain and 2 different pathway variants (see Figure 5F) were assayed. Bars show the mean. C) The difference in $^{13}$C-labeled fraction between $^{13}$C-formate-grown cells and $^{12}$C-formate-grown cells (mean and s.d., n=3).
